# Single molecule fingerprinting reveals different growth mechanisms in seed amplification assays for different polymorphs of αSynuclein fibrils

**DOI:** 10.1101/2024.03.05.583619

**Authors:** Derrick Lau, Yuan Tang, Vijaya Kenche, Thomas Copie, Daryan Kempe, Eve Jary, Noah J. Graves, Maté Biro, Colin L. Masters, Nicolas Dzamko, Yann Gambin, Emma Sierecki

**Author notes:** These authors contributed equally to this study (Derrick Lau, Yuan Tang).

## Abstract

Alpha-synuclein (αSyn) aggregates, detected in the biofluids of patients with Parkinson’s disease, have the ability to catalyze their own aggregation, leading to an increase in the number and size of aggregates. This self-templated amplification is used by newly developed assays to diagnose Parkinson’s disease and turned the presence of αSyn aggregates into a biomarker of the disease. It has become evident that αSyn can form fibrils with slightly different structures, called “strains” or polymorphs, but little is known about their differential reactivity in diagnostic assays. Here we compared the properties of two well-described αSyn polymorphs. Using single molecule techniques, we observed that one of the polymorphs had an increased tendency to undergo secondary nucleation and we showed that this could explain the differences of reactivity observed in *in vitro* seed amplification assay and cellular assays. Simulations and high-resolution microscopy suggest that a 100-fold difference in apparent rate of growth can be generated by a surprisingly low number of secondary nucleation “points” (1 every 2,000 monomers added by elongation). When both strains are present in the same seeded reaction, secondary nucleation displaces proportions dramatically and causes a single strain to dominate the reaction as the major end-product.

## INTRODUCTION

Alpha-synuclein (αSyn) aggregation is a hallmark of Parkinson’s disease (PD)^1^. Intracellular accumulation of aggregated αSyn in neurons is the main histopathological marker of PD and other synucleinopathies such as multiple system atrophy (MSA), and dementia with Lewy bodies (DLB). αSyn aggregates propagate from cell to cell, inducing protein aggregation in neighboring neurons in a prion-like manner^2^.

αSyn aggregation has been well-characterized *in vitro*^3^. αSyn monomers are mainly disordered in solution and aggregation happens *de novo* through misfolding. This leads to the formation of oligomers that will ultimately convert to cross-β-sheet structures that form the amyloid core of αSyn fibrils. These fibrils have the unique ability to propagate by recruiting monomers and incorporating them into the amyloid structure. As shown in Figure 1A, recruitment of monomers can occur at the end of the fibrils, causing linear growth of the fibrils in a mechanism called elongation. But recruitment of monomers can also occur along the surface of the fibrils and cause the formation of new fibrils growing as side branches; a mechanism termed secondary nucleation^4^.

**Figure 1.**
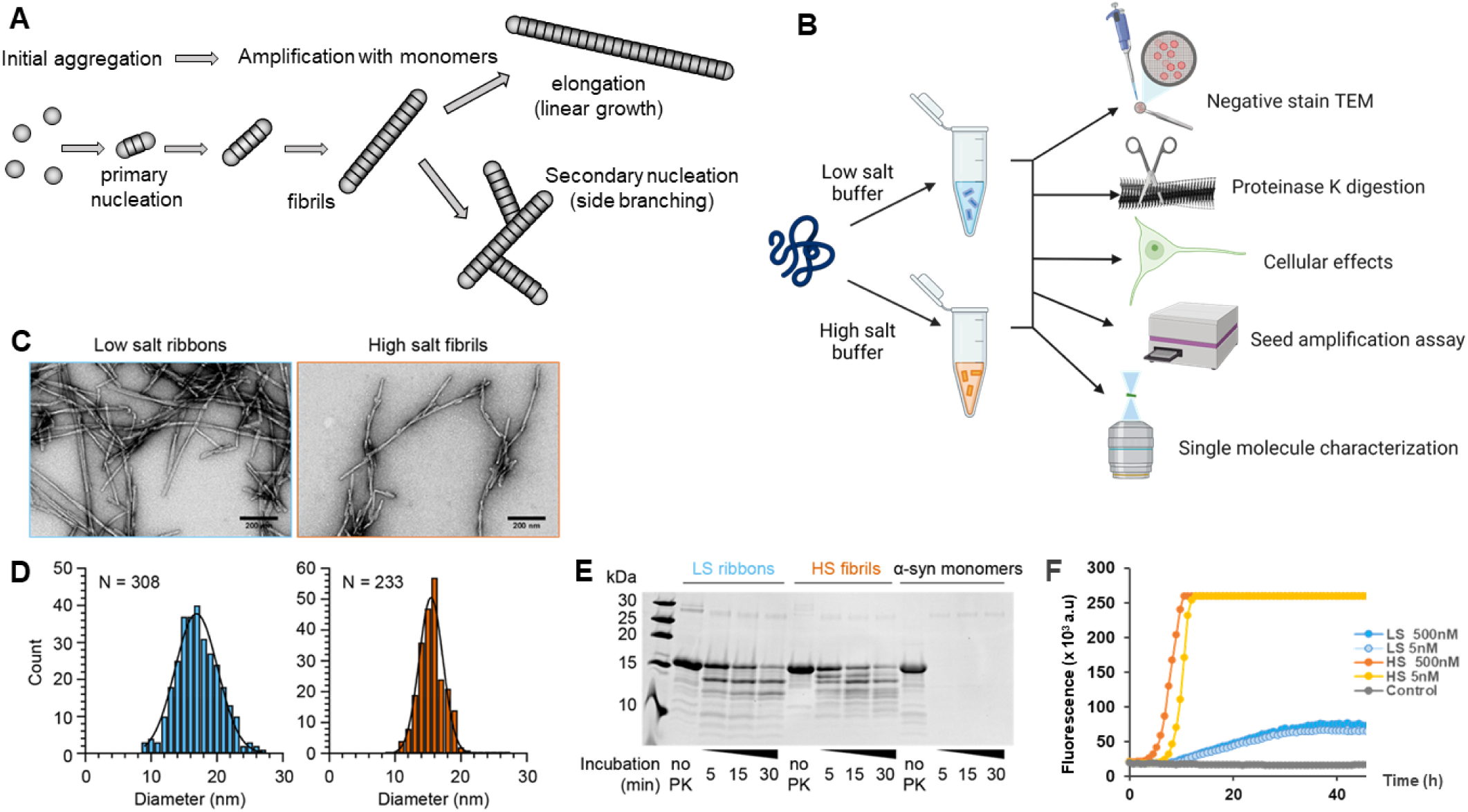
Characterization of *in vitro* recombinant αSyn assemblies. **A.** Schematic of αSyn aggregation and the different mechanisms at play: initial formation of aggregates by primary nucleation is followed by the growth of the fibrils by two mechanisms: pure elongation (linear growth) or secondary nucleation (creation of a new fibril on the surface of the first fibril). **B.** Schematic of the generation and characterization of recombinant αSyn polymorphs from monomers aggregated in low salt (sky blue) and high salt buffer (vermillion), using different assays. **C.** Transmission electron micrographs showing assemblies of recombinant αSyn under low salt and high salt buffer conditions, producing low salt ribbons and high salt fibrils respectively. Scale bar 200 nm. **D.** Histogram plot of the diameter of negatively stained low salt ribbons (17.1 ± 3.3 nm, mean ± standard deviation, N = 308) and high salt fibrils (15.5 ± 1.9 nm, mean ± standard deviation, N = 233) **E.** Proteinase K digestion profile of ribbons, fibrils and αSyn monomers. Substrates were digested by the addition of proteinase K (1.5 µg/mL), quenched at 5, 15, 30 min with loading buffer, heated at 90°C prior to resolving using reducing SDS-PAGE. **F.** Typical data obtained from a real-time quaking-induced conversion (RT-QuIC) assay on ribbons (LS) and fibrils (HS) at two different concentrations of seeds. Amplification was carried out in PBS, in the presence of 7 µM of WT αSyn monomer, 10 µM ThT and 6 glass beads. Fluorescence was recorded every 45 min. Control is an unseeded reaction.

This ability to convert monomer into amyloids in solution is exploited by the seed amplification assays (SAAs)^5^. SAAs were initially developed to detect misfolded cellular prion proteins (PrP^C^) in prion diseases and have been extended to identify the presence of other amyloids^6^. αSyn SAA is becoming a crucial diagnostic tool in PD research. A series of studies have demonstrated the sensitivity and specificity of αSyn SAAs to detect αSyn aggregates in the cerebrospinal fluid of patients and to accurately identify PD^5, 7, 8^. Positive responses to αSyn SAA correlated with an increased probability to develop a synucleinopathy disease, both for patients with REM sleep behavior disorder (RBD)^9, 10^ or mild cognitive impairment^11^ later developing PD and DLB, respectively. αSyn SAA was also recently applied to the study of a large longitudinal cohort of >1000 patients and found to be around 87.7% sensitive and >95% specific for PD^7^. Importantly, however, this study also revealed molecular heterogeneity between PD patients, with patients bearing a mutation in the PD risk gene leucine-rich repeat kinase 2 (*LRRK2*) showing less of an αSyn response than patients with sporadic PD. Taken together, these studies have established that αSyn aggregation is a diagnostic biomarker for PD and may allow for the earlier diagnosis of PD and other related synucleinopathies^7, 12, 13^.

Another important development in the field of PD and synucleinopathies is the emerging hypothesis that distinct αSyn strains characterize and perhaps dictate pathology^14–16^. The concept of strains was first established in prion research^17^ and refers to the fact that different prion polymorphs propagate specific disease phenotypes when injected into animal models. The concept has subsequently been extended to other amyloid-forming proteins, such as amyloid beta^18^, Tau^19^ and TDP-43^20^. In the case of αSyn, structural differences were observed in aggregates from the brain of PD/DLB^21^ patients compared to MSA patients^22, 23^ and there is growing evidence that αSyn strains induce different disease phenotypes^15, 16^.

αSyn strains (or polymorphs) can be generated *in vitro* by modifying the conditions in which αSyn aggregation occurs. Bousset and co-workers characterized two αSyn polymorphs, known as ribbons and fibrils^24^, by limited proteolysis and electron microscopy. These polymorphs were then shown to induce distinct toxicities in cells^24^. Intracerebral inoculation in rats showed that fibrils induced more neuronal death while ribbons induced more Lewy-body-like inclusions^25^. In wild-type mice, increased αSyn phosphorylation and ubiquitination was observed upon injection of ribbons into the striatum or olfactory bulb, compared to fibrils^26^. Increased αSyn deposition also correlated with ribbons injection in a mouse MSA model, while fibrils resulted in increased cellular death^27^. Finally, in non-human primates, intra-putaminal injection of ribbons and fibrils induced strain-specific patterns of αSyn aggregation and phosphorylation^28^.

Here we investigate the amplification propensity of these two polymorphs using single molecule spectroscopy^29, 30^ and different structural, biochemical and functional assays, as summarized in Figure 1B. We unexpectedly found that the mechanism of amplification differs dramatically between the two polymorphs, with the ribbons undergoing mainly elongation while the fibrils supported secondary nucleation. We show that this difference of amplification mechanism explains the differences of response in αSyn SAA. As SAA is becoming essential in PD diagnostics, the strain-dependent response and mechanisms of amplification should be taken into consideration in future studies.

## RESULTS

### Generation and characterization of ribbon and fibril polymorphs

Ribbons and fibrils were generated as previously described^15, 24^. Ribbons are obtained in low ionic conditions (5 mM Tris, pH 7.5, 0.02% w/v NaN_3_) while aggregation in high ionic conditions (50 mM Tris, pH 7.5, 150 mM KCl, 0.02% w/v NaN_3_) produced fibrils^24^. Well-established assays were performed to characterize these different polymorphs (Figure 1B).

Transmission electron microscopy revealed that monomeric αSyn effectively assembled to produce long filamentous structures after 7 days and 2 days of incubation for ribbons and fibrils, respectively (Figure 1C). Ribbons displayed a flattened morphology with occasional twists and had a measured diameter of 17.1 ± 3.3 nm (Figure 1C-D). Fibrils produce thinner cylindrical filaments with twists also observed, but they had a more consistent diameter of 15.5 ± 1.9 nm (Figure 1C-D). These measurements were consistent with previously published data^24^.

Proteinase K digestion was then performed to assess the profile of the ribbons and fibrils obtained. A time course of proteinase K digestion was performed for different batches of αSyn polymorphs. The β-sheet folded core of assembled αSyn filaments were highly resistant to proteinase while, as expected, monomers were completely digested after 5 min exposure to the enzyme. Digestion of the ribbons and fibrils revealed a distinct banding pattern, with more bands observed on digested fibrils compared to ribbons (Figure 1E). This demonstrates that these two assemblies were structurally different from each other. Notably, the banding patterns for our reference ribbons and fibrils were similar to those reported in Van der Perren et al^15^ indicating that the ribbons and fibrils were successfully recreated in the current study.

### Ribbons and fibrils have different profiles in SAA

We then tested the amplification potential of these two polymorphs in αSyn SAA. Seeded amplification was conducted in a real-time quaking-induced conversion (RT-QuIC) assay format, with intermittent shaking in a 96-well plate. Aggregation was monitored by following Thioflavin T (ThT) fluorescence over time, as ThT recognizes β-sheets structures aggregates.^31^ Ribbons or fibrils, at different concentrations were seeded in monomeric WT αSyn in PBS and shaken every 15 min at 300 rpm for 60 s. The kinetics were much faster for fibrils compared to ribbons (Figure 1F). We also observed that seeding with fibrils consistently led to a higher fluorescence plateau than ribbons at all concentrations tested. Therefore, fibrils are more efficient at templating aggregation, with ribbons and fibrils producing different profiles of amplification in RT-QuIC.

### Ribbons and fibrils induced distinct inclusions in SH-SY5Y cells

To evaluate the impact of the two different polymorphs on cells, differentiated SH-SY5Y neuroblastoma cells were treated with 5 μg/mL (0.35 μM) of ribbons or fibrils, or PBS as a control. The formation of intracellular αSyn inclusions was then measured over time at days 4, 7, and 10 post inoculation (Figure 2A). Both ribbon and fibril-treated cells exhibited visible inclusions and/or puncta of αSyn, while no such structures were observed in the control group (Figure 2B). Our previous results had shown that the number of αSyn inclusions reached the local maxima at days 4 to 6 post-fibril treatment before gradually declining, likely due to autophagic degradation^32^. Consistent with this, inclusions induced by both ribbons and fibrils reached their maximum on day 4 and steadily decreased subsequently on days 7 and 10 (Figure 2C). However, we observed significantly higher numbers of cytoplasmic inclusions induced by fibrils compared to ribbons at all three time points. Specifically on day 4, the intensity of αSyn inclusions induced by fibrils was approximately two-fold greater than those induced by ribbons, while by days 7 and 10, this difference had increased to more than four times. Meanwhile, cytotoxicity was not observed when we compared the extracellular release of lactate dehydrogenase between ribbon, fibrils treated cells and the control group (Supporting Figure 1). Note that the lack of cytotoxicity and cell death is in contrast to what Bousset et al.^24^ observed in their cell model.

**Figure 2.**
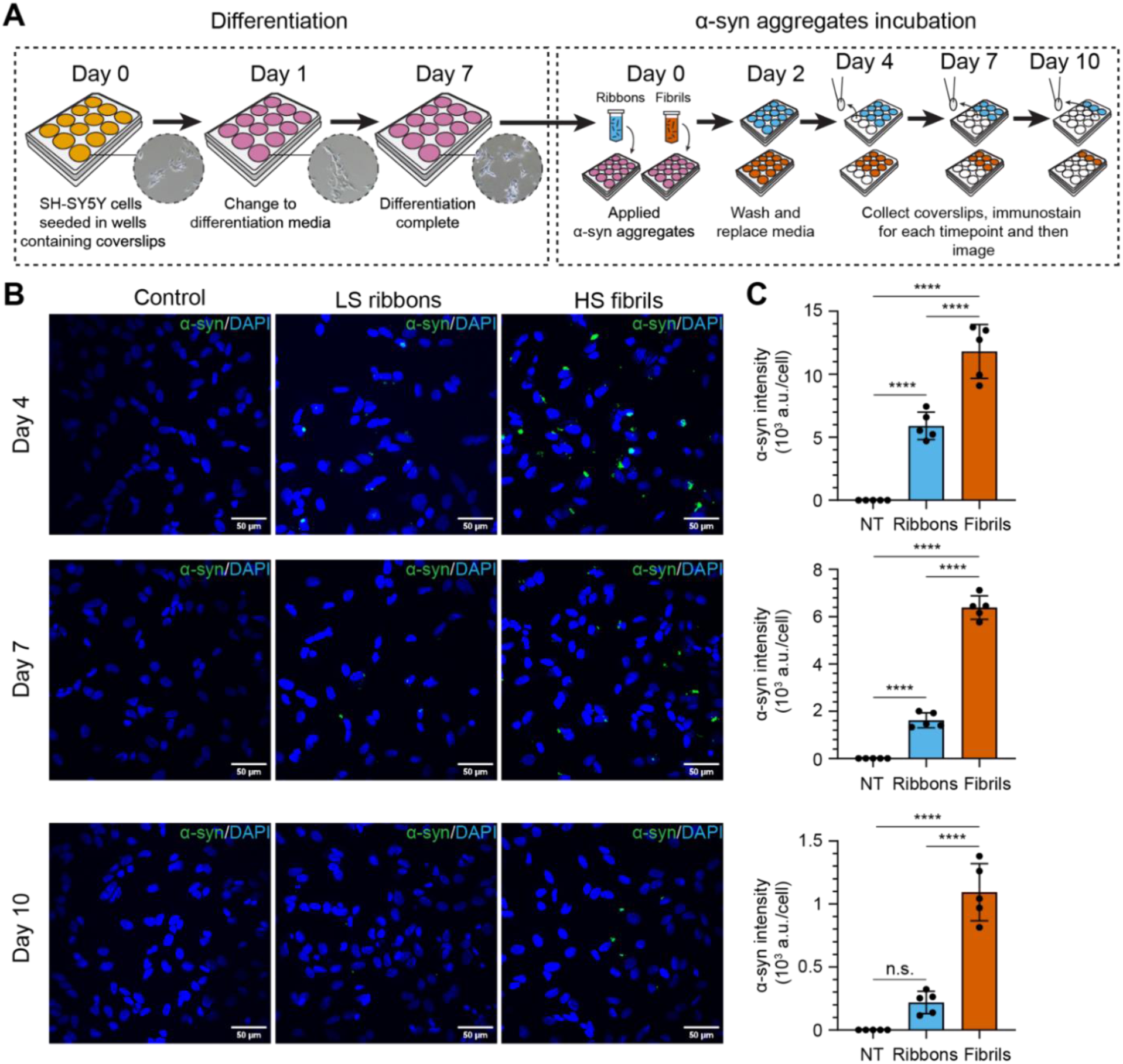
Characterization of recombinant αSyn assemblies incubation on SH-SY5Y cells. **A.** SH-SY5Y neuroblastoma cells culture scheme. Cells were seeded on coverslips at the bottom of 12-wells plate and imaged using brightfield microscopy (inset images). SH-SY5Y cells were differentiated for 7 days before incubation with αSyn ribbons or fibrils in differentiation media. Effects of αSyn assemblies on cells were assessed by immunostaining and confocal microscopy at 4, 7 and 10 days post addition of the assemblies. **B.** Differentiated SH-SY5Y cells were treated with or without 5 µg/mL of different αSyn polymorphs and subsequently fixed for immunofluorescence staining of total αSyn (green punctae) at indicated time points. DAPI staining is represented in blue. Scale bar is 50 μm. **C.** Bar graphs quantifying the average intensity of immunostained αSyn per cell. NT, no treatment, ribbons (sky blue) and fibrils (vermillion) respectively. Each symbol represents a FOV. Total αSyn intensity was normalized by the number of cells based on DAPI signal. Error bars represent mean ± standard deviation. Statistics used ordinary one-way ANOVA Tukey’s comparison test with a single pooled variance, *p* ≤ 0.0001 (****), nonsignificant (n.s.).

### Single molecule seed amplification assay revealed differential growth mechanism between ribbons and fibrils

Cell model data suggested that the high salt fibrils were more seeding competent with respect to their ability to generate bright aggregates in the cytosol and RT-QuIC data showed that fibrils amplify faster and are more abundant than ribbons. We then performed single molecule seed amplification assay (smSAA) measurements to examine the molecular properties of the amplified products, as previously described.^29, 30^ Sonicated seeds, either ribbons or fibrils, were diluted to single aggregate concentration in a solution of phosphate buffer saline (PBS) containing 10 µM ThT and 20 µM of monomeric human αSyn K23Q. We then characterized the aggregates before and after amplification (Figure 3A). As in RT-QuIC, the excess of monomeric αSyn promotes the growth of the fibrils in a prion-like manner. The main difference with RT-QuIC is that our protocol uses senescent conditions, as we want to avoid fragmentation to track the growth of the aggregates. In this work, we use a very short amplification time (5 h) as we have previously demonstrated that the technique remains sufficient to observe microscopic changes for individual aggregates.^29, 30^ To observe individual aggregates, we used our 3D-printed AttoBright instrument ^29, 30, 33^ as it was specifically designed to quantify and characterize ThT-positive events. The obtained fluorescence traces are real-time acquisition of the fluctuations of fluorescence in the small detection volume. As shown in Figure 3B, when a single aggregate diffuses through the detection volume, a burst of fluorescence above background is observed in the trace. These peaks are counted automatically using our algorithm and “fingerprinted”. For each peak, we determine its respective maximal intensity (prominence) and residence time (duration of the burst, which is linked to its physical size). The scatter plots of prominence against residence time provide a visualization of the different αSyn subspecies in the sample, as we introduced previously^30^. The area under the curve (AUC) for each peak is also calculated and the total intensity measured in all peaks of a given trace is an equivalent of the “bulk” readout of the RT-QuIC assay.

**Figure 3.**
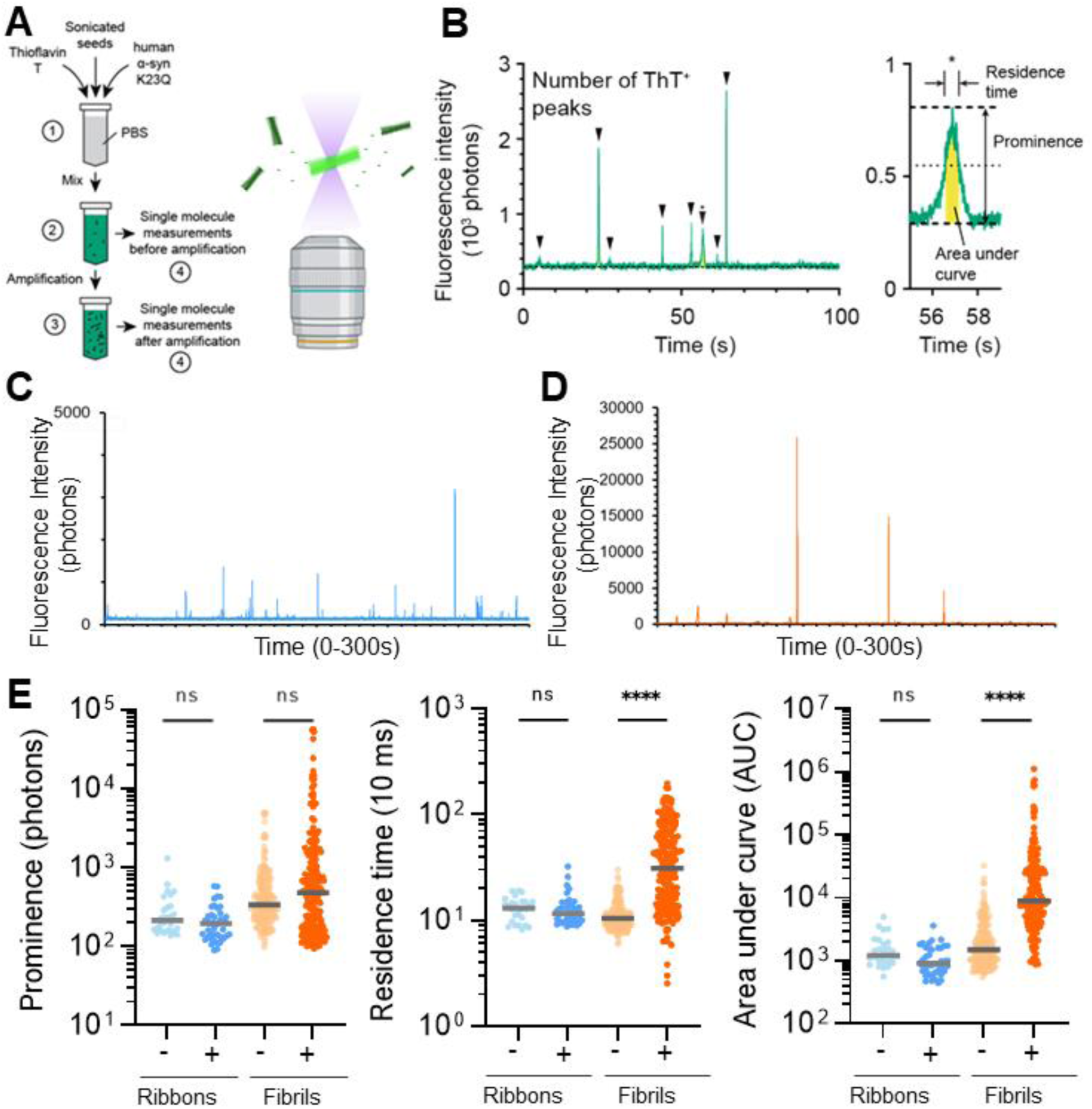
Single molecule measurement on recombinant αSyn polymorphs. **A.** Schematic of the single molecule seed amplification assay (smSAA). (1) αSyn polymorphs are seeded in a PBS solution containing thioflavin T (ThT, 10µM) and human αSyn K23Q (20 µM). (2) The solution is mixed, and half of the reaction is measured using single molecule spectroscopy to quantify the types of assemblies before an amplification step. (3) Sample is then amplified and measured again. Amplification enhances the signal of existing seeds by promoting αSyn assemblies. (4) Schematic of the microscope setup. The inset shows ThT stained αSyn assemblies diffusing across the confocal volume. **B.** Typical fluorescence traces. Fluorescence traces are analyzed to report the total number of ThT^+^ peaks (events) per measurement and (inset) the prominence of individual peaks, residence time (full width half-maximum), and area under the curve (yellow). The inset shows a region denoted by (*) in the trace. **C.** Typical raw trace obtained for amplified ribbons shows sharp peaks (fast diffusing) **D.** The amplified fibrils show a very different profile at the single molecule level, with very intense and slow diffusing species. **E.** Logarithmic scatter plot comparing peak prominence, residence time, and area under the curve before (light) and after amplification (dark) of LS ribbons and HS fibrils. Grey horizontal line is the median value. Each symbol represents an individual event and statistics used Kruskall-Wallis test, *p* ≤ 0.0001 (****), nonsignificant (n.s.).

Before amplification, ribbons and fibrils were sonicated to reduce the size of the seeds which was then confirmed by electron microscopy (Supporting Figure 2). The single molecule fluorescent traces show that before amplification, both ribbons and fibrils appear as fast diffusing aggregates. After only 5 h of amplification, ribbons still behave as fast diffusing species (Figure 3C) while we observed a very large change in the properties of fibrils, as the raw traces show very wide and intense peaks (Figure 3D). The fingerprinting of the intensity, residence time and area under the curve for the ribbons and fibrils show two very distinct signatures (Figure 3E). The most differentiating feature is the large increase in residence time for the fibrils, and this suggests two very different mechanisms of growth for the two polymorphs.

### High salt fibrils grew by secondary nucleation

The large shift in residence time observed for amplified fibrils reveals that their apparent size has greatly increased. We therefore turned to high-resolution microscopy to observe these large aggregates. Confocal microscopy images show that within 5 h, nanometer-sized sonicated fibrils have grown to sizes up to 5 microns (Figure 4). The 3D reconstruction of the aggregates clearly shows the growth of dense networks of fibrils, with branching structures and not pure linear elongation of a long and single amyloid core (Supporting Figure 3). This aligns perfectly with the idea of secondary nucleation and fibril-growth catalyzed on the surface of the fibrils.

**Figure 4.**
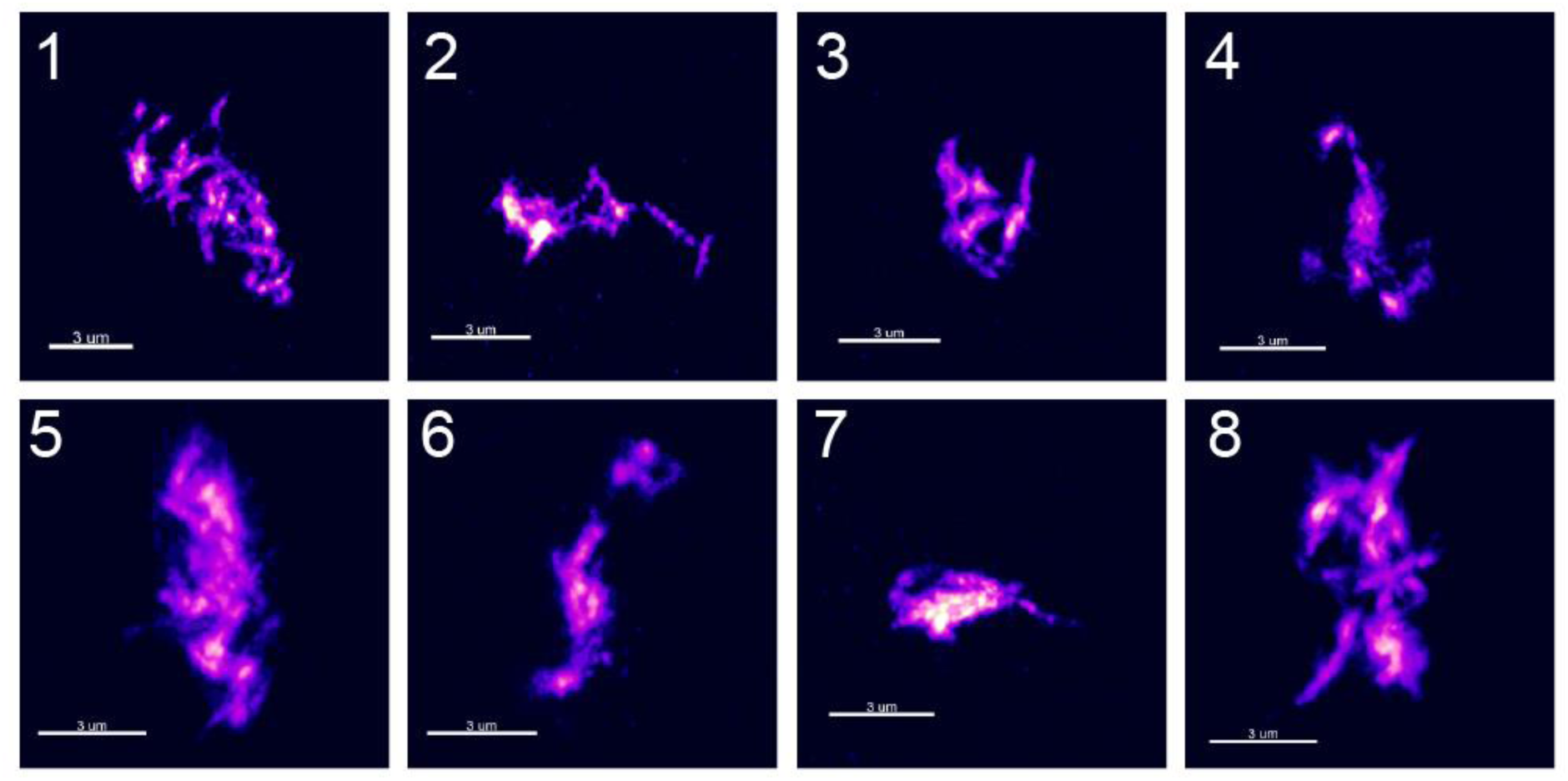
high-resolution confocal imaging of the amplified fibrils. Confocal microscopy was performed on the fibrils after 5 h of amplification, on a Zeiss LSM 880 microscope with an AiryScan unit (see SI for details). Z-stacks were recorded and analyzed with Zen software to generate the final high-resolution images. In the same setup, amplified ribbons could not be detected, as expected for smaller, fast-diffusing particles.

It should be noted that in general, secondary nucleation is proposed to release new fibrils into solution, but our images suggest that the new fibrils do not detach spontaneously. Confirming this idea, the number of peaks measured by the single molecule technique did not change significantly during amplification. This could be easily explained by our amplification conditions, as the solution is not shaken, and we do not use any mechanical disruption (beads or sonication) that could break the assemblies.

### Impact of secondary nucleation on seed amplification assays

Having observed that fibrils amplify much faster than ribbons, we wanted to see if those differences could be explained by the addition of secondary nucleation and determine what rate of secondary nucleation would be sufficient to dominate the kinetics. We turned to mathematical modelling following the approach described in Knowles et al^34^. In this work, the authors had laid the foundations for a global fit of the kinetics found in seed amplification assays.

We considered three elementary reactions in the model: (1) **elongation** - adding a monomer on a growth point, (2) **secondary nucleation** - adding a new growth point in a polymer, and (3) **fragmentation** - breaking a polymer in two and creating two new growth points. Those three processes are schematized in Figure 5A. In this model, we did not consider primary nucleation as seed amplification assays are designed to eliminate spontaneous formation of seeding competent αSyn nuclei in the absence of aggregates. The detailed equations can be found in the supplementary information section.

**Figure 5.**
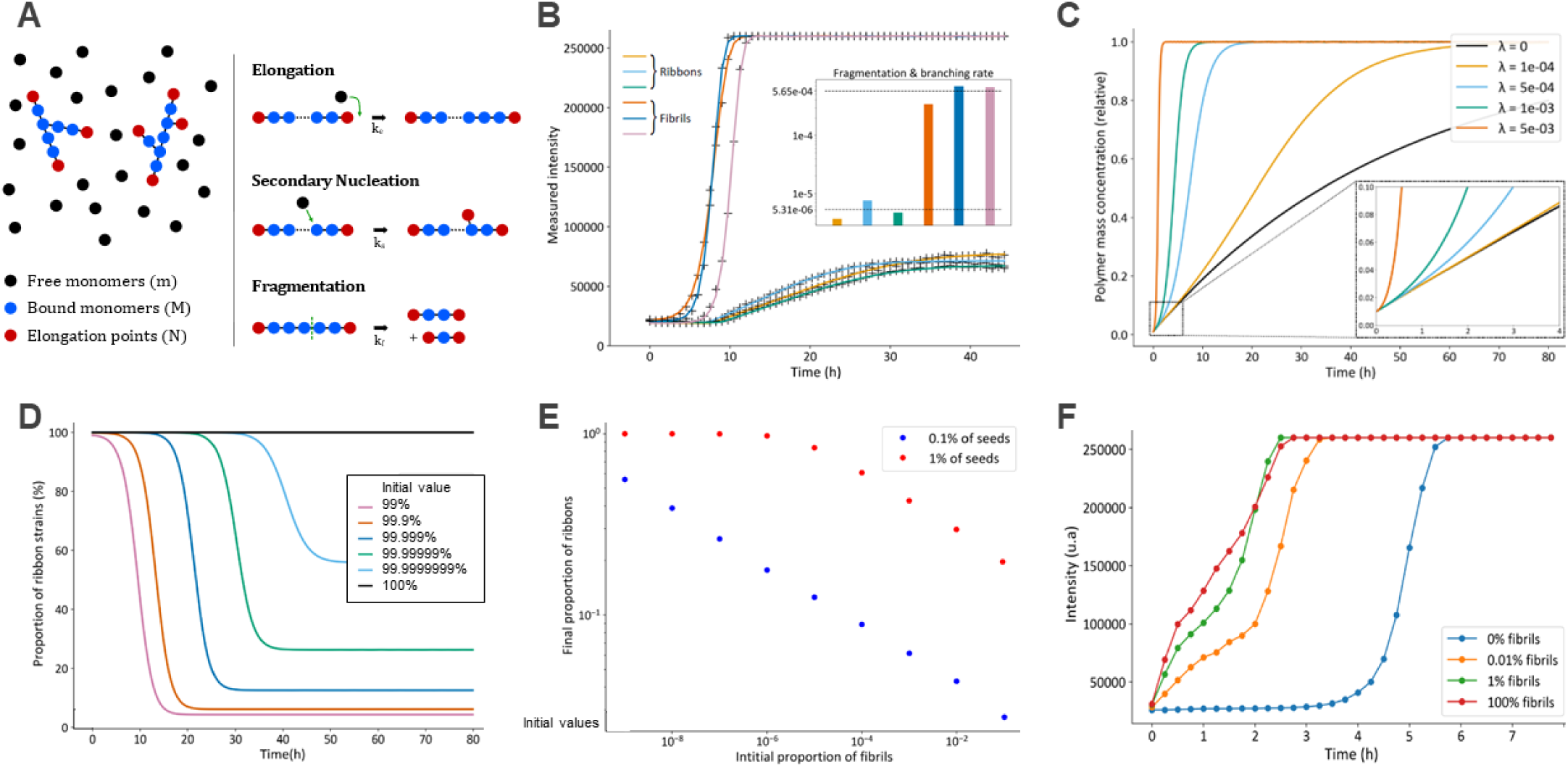
Mathematical modeling of seed amplification and competition between strains. **A.** Visual representation of the different parameters used for simulation: *M* - the polymer mass concentration, *N* - the concentration of growth points, and *m* - the concentration of free monomers. The diagram on the right shows the elementary reactions that we considered in this model. *k_e_* (resp. *k_s_*, *k_f_*) is the rate of the elongation (resp. secondary nucleation, fragmentation) reaction. **B.** Numerical fit of experimental RT-QuIC data seeded with ribbons or fibrils at 5, 50 or 500 nM using the numerical solution derived from panel A. **C**. Numerical solutions for the time evolution of the relative polymer mass concentration, with different secondary nucleation to elongation ratios denoted by λ where λ = 10^-3^ means that one new branch is created for 1000 monomers added through elongation. The relative polymer mass reaches 1 when all monomers have been incorporated in the polymers. Fragmentation rate is set to 0. **D.** Numerical modeling of competition between different strains seeded in the same solution. Fragmentation and elongation rates were set to the same value, and a rate of secondary nucleation was applied only for fibrils. The rate constants were extracted from the fit in panel B. The initial composition of polymer mass were set to 100%, 99.9999999%, 99.99999%, 99.999%, 99.9% and 99% of ribbons. **E.** Final content of ribbons in the polymer mass at the end of simulation (80 h) is displayed for low seed concentration (0.1% of monomer equivalent) or high seed concentration (1% monomer equivalent). **F.** Experimental competition between ribbons and fibrils. The 4 curves correspond to 4 different ratios of ribbons and fibrils in the seeds. Each curve is obtained by averaging the results of 2 RT-QuIC experiments with the same parameters.

We simulated the RT-QuIC responses obtained for ribbons and fibrils, using the same elongation rate for the two species and fitted the experimental curves by adjusting a combined rate of fragmentation and secondary nucleation 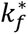. As shown in Figure 5B, the values obtained for ribbons 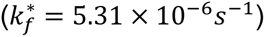 and fibrils 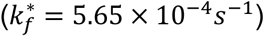 differ by 100-fold. As the samples in the 96 wells plate are shaking at the same speed with the same number of beads, it would be surprising that the rate of fragmentation would be extremely different, and we can assume that a large contribution to the kinetics come from the presence of secondary nucleation.

To go further, we decided to simulate the effect of secondary nucleation and generated growth curves for different values of k_s_ and k_e_. To simplify, we consider the parameter 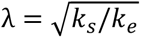 that describes the number of branching events. For example, λ = 10^-4^ means that approximately one point of secondary nucleation is created every 10,000 elongation events. The curves are presented in Figure 5C. The insert demonstrates that for λ = 10^-4^, the effect of secondary nucleation is barely visible at short time scales, but significant deviations are observed for λ = 5x10^-4^. With λ = 10^-3^ i.e. one nucleation every 1000 monomers, the apparent growth doubled within 2 h, and the reaction proceed extremely rapidly.

These simulations match the data observed for fibrils and ribbons and enabled us to estimate the number of secondary nucleation events. The equations suggest that the 100-fold difference in rate of growth can be generated by as little as one secondary nucleation “point” every 2,000 monomers added by elongation (see Supplementary Information for details of these estimations.). Considering a typical interstrand distance of 0.5 nm in amyloid fibrils, this suggests that branching would occur every 1 µm; the 3D images of the “specks” after amplification shown in Figure 4 are consistent with this spacing between secondary branches.

Such a large difference in kinetics prompted us to explore competition between strains and see whether there is selection of specific strains during amplification. Indeed, recent publications have shown that different αSyn strains are produced in the different synucleinopathies. The question now is whether the assays can replicate the seeds with high fidelity, or if they can be strongly biased towards strains with high secondary nucleation propensity.

We first simulated amplification of samples where ribbons and fibrils would be present simultaneously. We considered a constant total number of “seeds” (ribbons + fibrils) and only change the ratio of the two polymorphs. We simulated growth of the seeds and plotted the proportion of ribbons as a function of time. Figure 5D shows that extremely small amounts of fibrils present in solution would dominate the response. The effect is so significant that even with an initial proportion of ribbons close to 100% (99.99999%), fibrils become the dominant polymorph at the end of the reaction.

It is interesting to note that when seeds are present in high numbers (up to 1% of seeds) the replication has higher fidelity (Figure 5E). This is relevant for polymorph propagation *in vitro* (when re-amplifying a stock of highly concentrated fibrils) but in seed amplification assays from biofluids, the initial concentration of aggregates is likely to be in the picomolar range (i.e. 10^6^ lower concentrations than the monomer used for amplification.)

Finally, we tested the competition between polymorphs experimentally, by setting up RT-QuIC reactions with different ratios of fibrils to ribbons. The data obtained in Figure 5F confirm that small amounts of fibrils (as low as 0.01% of ribbons) dramatically change the overall response of the RT-QuIC assays.

### Assessment of growth mechanisms under different experimental conditions

The mechanism of amyloid growth is known to be highly dependent on experimental conditions and we wanted to examine whether the difference between fibrils and ribbons amplification was conserved in other amplification buffers. We next performed smSAA using different protocols and assessed their single molecule profiles.

First, we performed smSAA for 5 h using different buffer conditions outlined in Figure 6A. These conditions were selected based on their use across different labs to perform RT-QuIC on patient samples.^5^ Amplification (measured as the increase in total ThT fluorescence) occurred in most conditions (Figure 6B), and again a strong difference was observed between ribbons and fibrils for efficiency of growth. Fibrils were amplified strongly in all buffers, with at least 7-fold increase in total ThT fluorescence per trace (Figure 6B). As with our reference conditions (condition D, PBS, K23Q αSyn monomer), all conditions created a significant increase in residence time (Figure 6C). Ribbons on the other hand generally failed to increase in residence time but rather showed higher prominence. One notable exception is condition B. Condition B provided the best conditions to promote the growth of ribbons, with an 8.5-fold change in total fluorescence per trace. Notably, this condition was the only one that led to the appearance of high molecular weight αSyn “ribbons” which shifted the median residence time from 116 to 157 ms (Figure 6C). The distinctive buffer constituents of B include its mild acidic conditions (pH 6.5) coupled with high ionic strength (500 mM NaCl). Acidic conditions have been shown to accelerate aggregation of αSyn due to enhancement of secondary nucleation^4,35^. The effect of the high concentration of NaCl could enhance the rate of aggregation because the high salt buffer produced significantly more yield and faster assembly kinetics compared to ribbons.^24^ However, some studies demonstrated that very high ionic strength significantly reduces the aggregation of αSyn due screening of electrostatic repulsion by salt^36, 37^. Condition B also produced the largest change in median residence time for fibrils, from 123 ms to 628 ms. These data strongly suggest that the residence time can be used as a signature of secondary nucleation.

**Figure 6.**
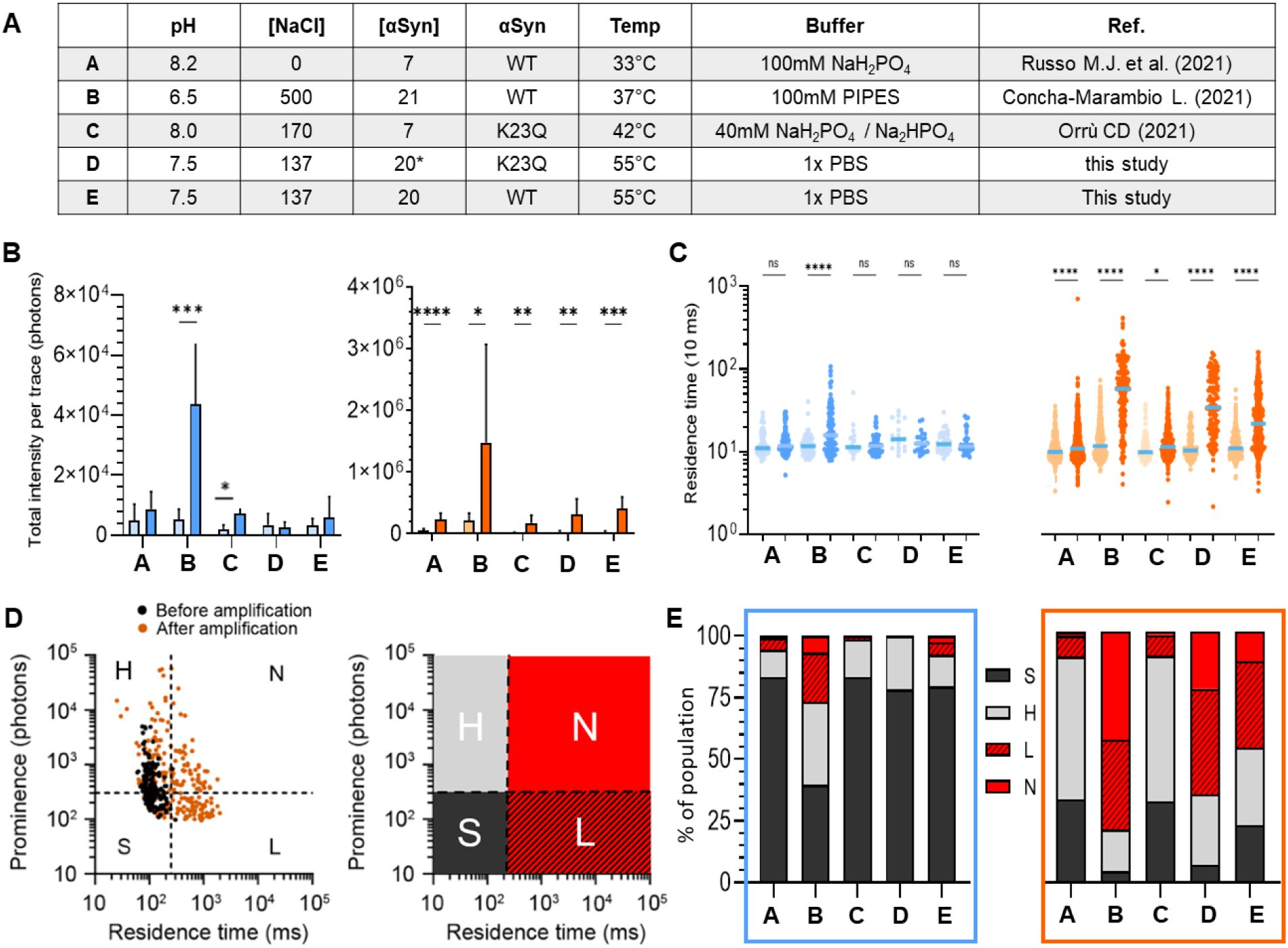
Single molecule profiling of ribbons and fibrils amplified in different conditions. **A.** Table of the different conditions used: A (32°C, 7 µM αSyn WT), B (37°C, 21 µM αSyn WT), (42°C, 7 µM αSyn K23Q), D (55°C, 20 µM αSyn K23Q), E (55°C, 20 µM αSyn WT). **B.** Bar graph showing the total fluorescence intensity if the trace for low salt ribbons (blue) and high salt fibrils (vermillion) before (light) and after 5 h amplification (bright) in the different conditions. Error bar is standard deviation. **C.** Logarithmic scatter plot comparing the residence time of ThT^+^ peaks (events) detected in fluorescence traces before (light) and after amplification (bright), for ribbons (blue) and fibrils (vermillion). Each symbol represents an individual event and statistics used Kruskall-Wallis test, *p* ≤ 0.05 (*), *p* ≤ 0.0001 (****), nonsignificant (n.s.). **D.** The events before (black) and after (vermillion) amplification (here for fibrils amplified in conditions D) can be visualized on a scatterplot of prominence vs residence time. We defined 4 quadrants to separate fast-diffusing and slow diffusing species, and high-intensity vs low-intensity species. The four species are defined as S (dark grey, < 300 photons, < 250 ms), H (light gray, > 300 photons, < 250 ms), L (hashed red, < 300 photons, > 250 ms) and N (red, > 300 photons, > 250 ms). **E.** Distribution of the different species detected after amplification for the different buffers. Results for the ribbons and fibrils are presented in the blue and vermillion frame, respectively.

We then analyzed the “fingerprints” of the aggregates by plotting the intensity of the peak vs. residence time, similarly to FACS (fluorescence-activated cell sorting). We “gated” at different thresholds of intensity and residence time to define 4 sub-species, as shown in Figure 6D. For ribbons, the relative low abundance of slow diffusing particles (L and N quadrants) indicates that growth mainly occurs through elongation except in acidic conditions where secondary nucleation could be observed. Fibrils produced large assemblies (L and N) when amplified in PBS WT and K23Q (conditions D and E), similarly to B, indicating a predominance of secondary nucleation. We also observed that conditions A and C produced a similar profile with a majority of H events, suggesting that these two conditions partially suppressed secondary nucleation while promoting elongation to maintain the strong amplification of fibrils (Figure 6E). These experiments are performed in mildly basic buffer (phosphate buffer, pH ≥ 8.0) which is indeed thought to limit secondary nucleation^4^.

We next examined the effects of other experimental parameters on the mechanism of amplification. Changing the amplification conditions from 3 h at 55°C to 5 h at 37°C did not significantly change the efficiency or mechanism of growth (data not shown). SmSAA was then performed in PBS with various concentrations of αSyn WT to determine the minimum concentration required to observe secondary nucleation. With ribbons, some increase in total ThT fluorescence was observed with substrate concentration ≥ 7 μM but all conditions failed to increase the median residence time after amplification, even at the highest concentration of 28 μM monomer (Supporting Figure 4). Fibrils had an observable change in total ThT fluorescence at 3.5 μM of monomers, emphasizing their stronger seeding potential. However, changes in residence time were observed as the basis of the growth mechanism only when ≥ 7 μM of monomers were supplied in the reaction.

We then paid particular interest to the effect of salt concentration on amplification (Figure 7). Contrary to other amyloids, it was reported that increased salt concentration reduced the kinetics of amplification of αSyn fibrils^37^ ^36^. We performed two series of RT-QuIC and smSAA experiments, varying the salt (NaCl) concentration from 0-500 mM in neutral (phosphate buffer, pH 7.4) and acidic (MES, pH 6.0) conditions. In Figure 7A and 7B, the results of the RT-QuIC are shown for amplification of the fibrils in different salt concentrations at low pH and neutral pH. In neutral pH, increasing salt concentration leads to faster reactions (Figure 7A) while we observe the reverse effect for low pH (Figure 7B). The RT-QuIC curves can be quantified by measuring the slope of the reaction, the time to initial threshold (TTT), and time to reach mid-reaction (TTM), as described in Figure 7C. We then conducted smSAA on fibrils and ribbons in these conditions and studied correlations between the smSAA and RT-QuIC. To take into account the faster dynamics of secondary nucleation, single molecule traces were acquired after 1.5 h and 4.5 h, for acidic and neutral conditions respectively.

**Figure 7.**
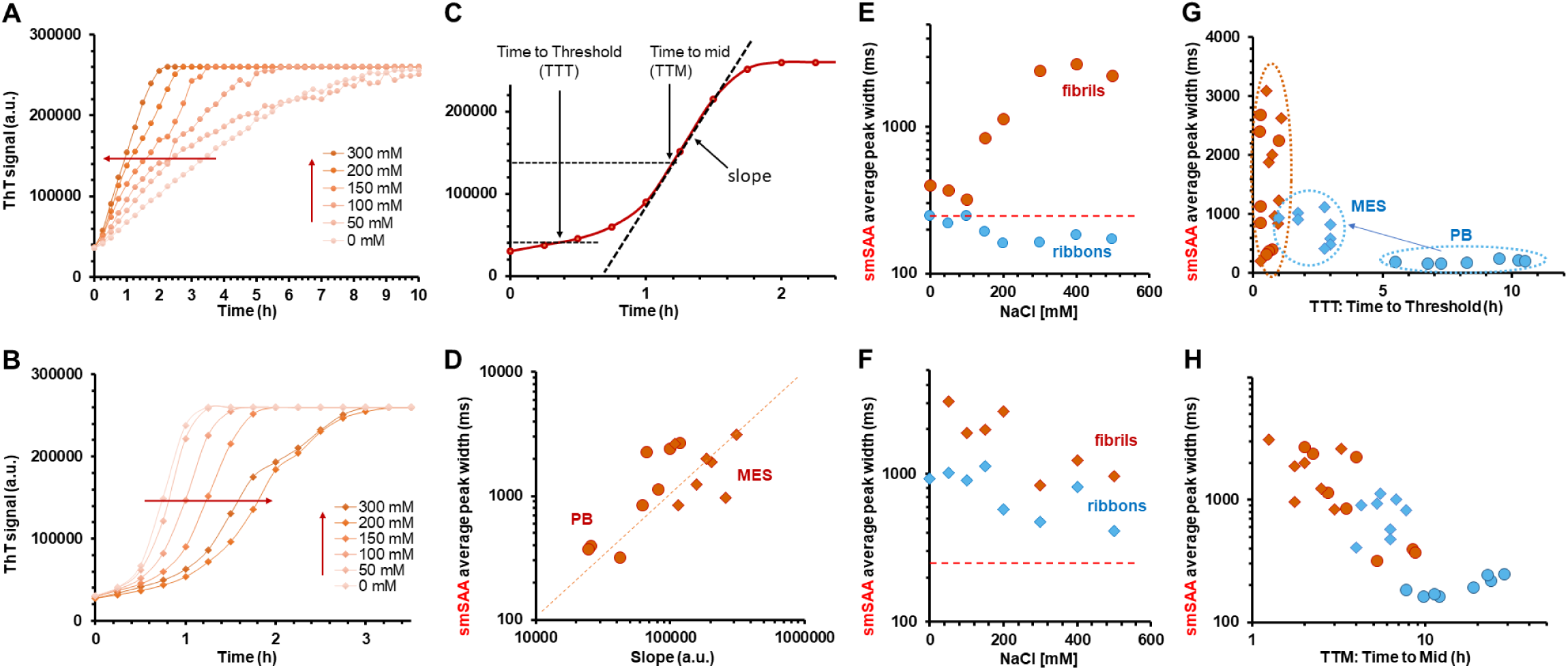
Correlation between RT-QuIC and single molecule profiling and effect of salt concentration. **A.** RT-QuIC curves obtained for reactions seeded with fibrils in Phosphate Buffer (PB, pH 7.5) with different concentrations of NaCl. The curves show an increase in reactivity with increasing salt concentrations. **B.** RT-QuIC curves obtained for reactions seeded with fibrils in MES buffer (pH 6.0) with different NaCl concentrations; the reactions appear slower when [NaCl] increases. **C.** Schematic of the RT-QuIC curve and definition of different parameters for quantification: slope of the reaction (calculated around the mid-point), time-to-threshold (TTT, timepoint when reactivity reaches 10% of the final value), and time-to-midpoint (TTM, timepoint when ThT signal has increased to 50% of the final value). **D.** From experiments conducted in quiescent conditions and analysed using our single molecule method, we determined the smSAA average peak width. The peak width correlates strongly with the slope of the RT-QuIC reaction, as shown here for fibrils in PB (round) and MES (diamond) buffers. Data for ribbons are color-coded in blue, and in red for fibrils; data are displayed as round dots for Phosphate Buffer, and diamonds for MES buffer. These colours are used for panel E-H. **E.** smSAA diffusion time as a function of salt concentration for ribbons (blue) and fibrils (vermillion) in Phosphate Buffer pH 7.5. The red dotted line indicates the arbitrary threshold for slow diffusing particles (250 ms). **F.** SmSAA diffusion times for ribbons (blue) and fibrils (vermillion) in MES Buffer pH 6.0; the data show that both fibrils and ribbons are produced slow diffusing species. **G.** Scatter plot of smSAA average peak width vs Time to Threshold (TTT), showing that low pH leads to much faster reactions for ribbons and higher diffusion times. **H.** Scatter plot of Time to Mid (TTM) vs smSAA average peak width for ribbons and fibrils in all conditions of pH and salt, showing a general trend of decreasing diffusion times with increasing reaction times.

As shown in Figure 7D, the slope of the RT-QuIC reactions depends very strongly on the diffusion times measured in smSAA. When we explore into more details the salt-dependence of the diffusion times, we observe two different behaviors. At neutral pH, fibrils amplified significantly better than ribbons, as observed before; we observe a clear salt-dependent increase in the diffusion times, suggesting that secondary nucleation is increasing as a function of salt concentration (Figure 7E). For low pH, we observe higher average diffusion times for both ribbons and fibrils, and a decrease of diffusion times with increasing salt concentrations (Figure 7F). As shown in Figure 7G, the switch to low pH increases the reactivity of ribbons, as we observe a shift to lower time to threshold for all salt conditions; at the same time, we detect a large shift in diffusion times. This suggests that acidic conditions could boost the apparent RT-QuIC reactivity of ribbons by increasing secondary nucleation. This matches well with published observations^4^. In Figure 7H, the overall data for ribbons and fibrils, in low and neutral pH, show a clear trend of faster reactions with higher smSAA diffusion times, reinforcing the idea that secondary nucleation plays a major role in the apparent reactivity in RT-QuiC.

### αSyn polymorphs maintained their properties upon amplification

Previous results by Bousset et al.^24^ demonstrated that the low salt ribbons and high salt fibrils gave rise to different phenotypes when seeded in different buffers, pointing to a loss of properties upon amplification. To re-explore this aspect, ribbons were seeded in a high salt buffer and fibrils were seeded in a low salt buffer in a cross-buffer seeding experiment to produce new assemblies. The polymorphs generated this way were denoted as LSxH and HSxL respectively. Ribbons seeded in low buffer (LSxL) and fibrils in high buffer (HSxH) were also employed as controls. The cross-seeded species were assessed by transmission electron microscopy, proteinase K digestion and smSAA. LSxL retained the same phenotype as its parental ribbon seeds, and no variation was observed between two independent batches LSxL-1 and LSxL-2 (Figure 8B-C). These products produce filaments of similar width, had identical proteinase K digestion pattern and identical single molecule amplification phenotype in its poor ability to undergo secondary nucleation to its parental strain as expected (Figure 8B-E). LSxH also resembles its parental LS strain and does not have the same PK digestion profile as fibrils, which is recognized by a discriminatory band at approximately 13 kDa (Figure 8C, black arrow). This indicates that free monomers do not sample the high ionic buffer to generate high salt fibrils *de novo* when ribbons seeds are present in solution. Surprisingly, cross-seeded HSxL also retained the PK profile and secondary nucleation prone phenotype in smSAA of the parent fibrils. HSxL was indeed also highly seeding competent and able to grow via secondary nucleation (Figure 8D-E). HSxL PK digestion did not reveal any new bands of *de novo* assembly of low salt ribbons in solution. These findings indicate that the amyloid core, at least with the recombinant *in vitro* polymorphs used in the current study, can propagate structural information irrespective of the environment in which they are amplified.

**Figure 8.**
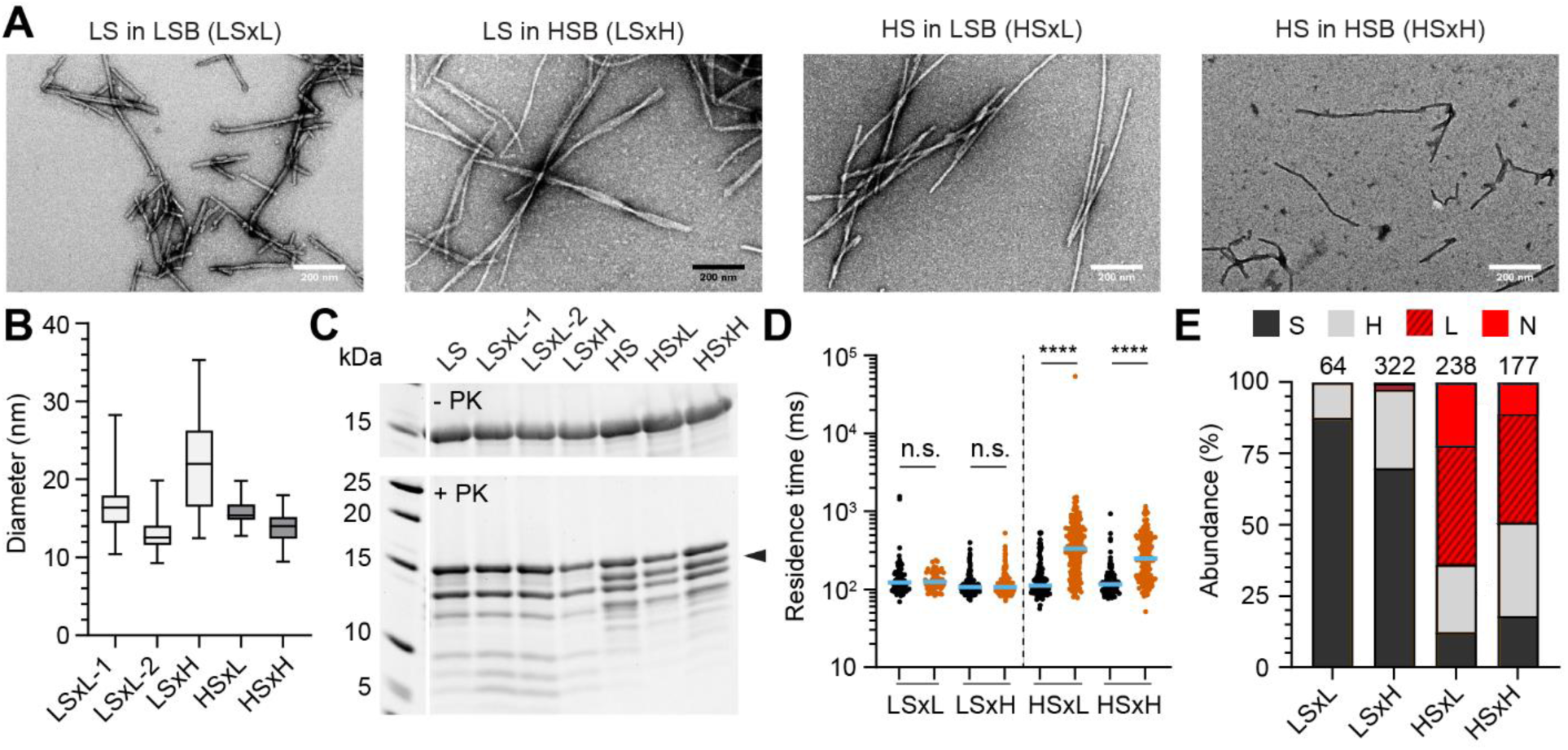
Characterization of αSyn polymorphs obtained by cross-buffer seeded reactions. **A.** Ribbons (LS) were seeded in low salt buffer (LSB) or high salt buffer (HSB) yielding the products LSxL and LSxH, respectively. Similarly, fibrils (HS) were seeded in low salt buffer (LSB) or high salt buffer (HSB) giving rise to HSxL and HSxH products. Representative electron micrographs of negatively stained assemblies after amplification. **B.** Box plot quantifying the diameter of filaments from electron micrographs; LSxL-1 (16.4 ± 2.7 nm, mean ± standard deviation, N = 244), LSxL-2 (13.0 ± 2.2 nm, N = 42), LSxH (21.9 ± 6.0 nm, N = 40), HSxL (15.8 ± 1.6 nm, N = 76), HSxH (13.7 ± 2.0, N = 28). Whiskers represent min-max values. **C.** Proteinase K digestion profile of seeded products from recombinant αSyn polymorphs in the absence (top) and after addition of proteinase K (bottom, PK). Black arrow points at a band that discriminates ribbons from fibrils. **D.** Logarithmic scatter plot comparing the residence time of events in seeded reaction before (black) and after (vermillion) amplification with αSyn K23Q. The median value is highlighted in blue. Each symbol represents an individual event and statistics used Kruskall-Wallis test, *p* ≤ 0.0001 (****), nonsignificant (n.s.). **E.** Distribution of ThT^+^ events detected after amplification in cross-seeded reactions. S (dark grey, < 300 photons, < 250 ms), H (light grey, > 300 photons, < 250 ms), L (hashed red, < 300 photons, > 250 ms) and N (red, > 300 photons, > 250 ms). The number of events detected and used for quantification is indicated above each bar.

As a final validation strategy, we sought to explore the potential variations of the pathological properties of these cross-seeded polymorphs in SH-SY5Y neuroblastoma cells. Large batches of cross-buffer seeded polymorphs LSxL, HSxH, LSxH and HSxL were prepared and characterized by single molecule amplification and proteinase K digestion (Supporting Figure 5). These cross-seeded polymorphs were added to differentiated SH-SY5Y cells to assess their ability to induce αSyn inclusions in the cytosol using immunostaining and confocal microscopy. The density, count, and intensity of αSyn of LSxH were not significantly different from cells treated with LSxL (Figure 9A-B). LSxH produced low αSyn intensities per cell, similar to the parental strain (Figure 9B). Inversely, HSxL were able to produce very bright and large αSyn punctae when seeded in cells, indistinguishable from HSxH (Figure 9A-B). Notably, it was found that cells exposed to both fibrils HSxL or HSxH exhibited significantly stronger cytoplasmic αSyn intensity than those exposed to ribbons on day 4, 7, or 10. The cross-buffer seeded polymorphs were not cytotoxic to the cells, similar to the parent strains (Supporting Figure 6). Overall, these results were consistent with the previous data (Figure 8), as there was no disparity in cytoplasmic inclusions between ribbons or fibrils seeded in their parental buffer or cross-seeded in high salt or low salt buffer respectively^24^.

**Figure 9.**
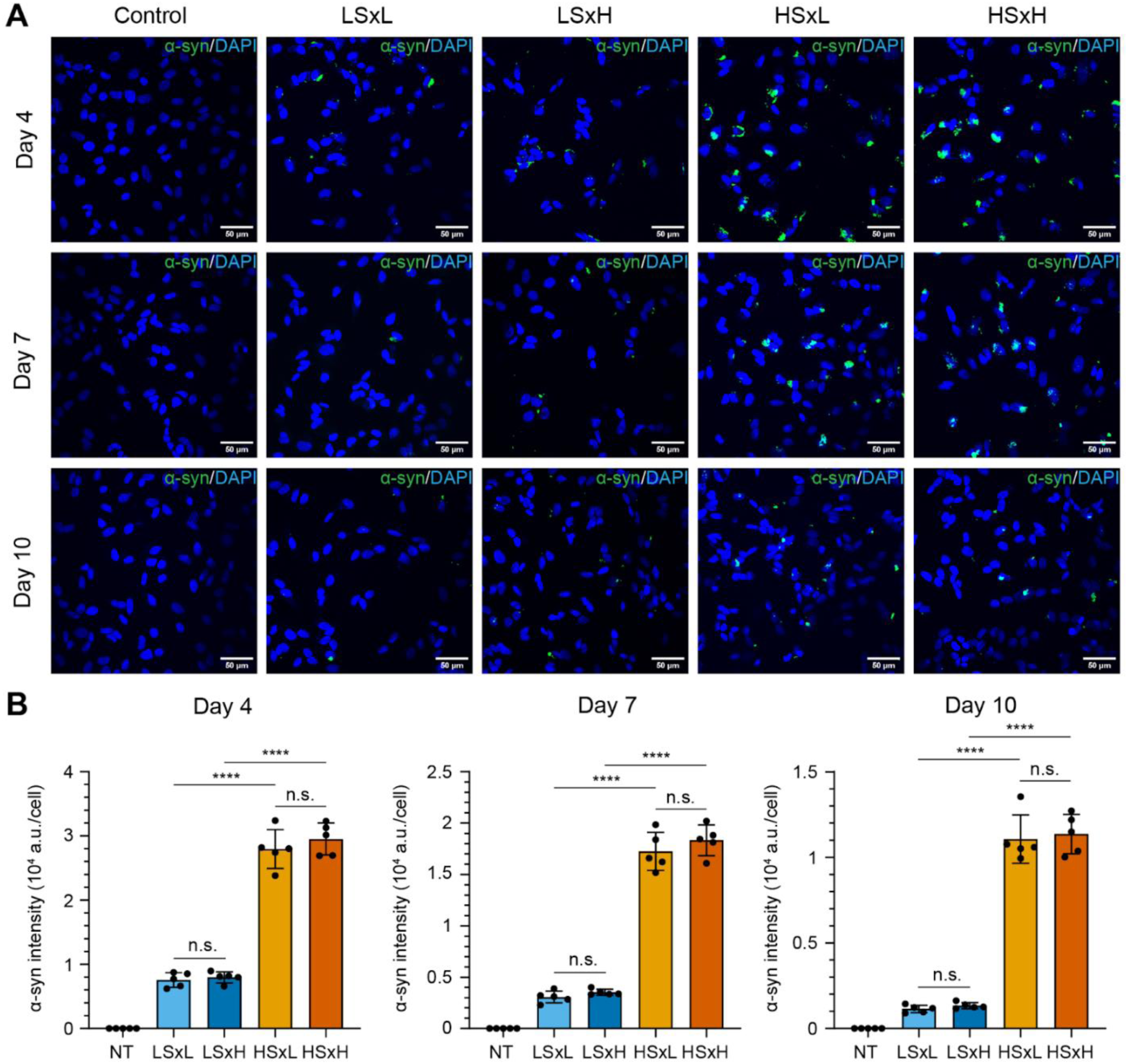
Temporal changes of cytoplasmic αSyn inclusions following treatment of SH-SY5Y cells with cross-seeded αSyn aggregates. **A.** Confocal images of differentiated SH-SY5Y cells treated without (NT, no treatment) or with 5 µg/mL of diverse αSyn aggregates: ribbons seeded in low salt buffer (LSxL), ribbons seeded in high salt buffer (LSxH), fibrils in low salt buffer (HSxL) and fibrils seeded in high salt buffer (HSxH). Cells were subsequently fixed for immunofluorescence staining to quantify total αSyn (green) at day 4, 7 and 10 post incubation with the aggregates. DAPI staining is represented in blue. Scale bar of 50 µm. **B.** Bar graphs quantifying the average intensity of immunostained αSyn per cell upon treatment with different αSyn aggregates (NT: not treated, light blue: LSxL, dark blue: LSxH, light vermillion : HSxL, dark vermillion: HSxH). Each symbol represents a FOV. Total αSyn intensity was normalized by the number of cells based on DAPI signal. Error bars represent mean ± standard deviation. Statistics used ordinary one-way ANOVA Tukey’s comparison test with a single pooled variance, *p* ≤ 0.0001 (****), nonsignificant (n.s.).

## DISCUSSION

### Generating fibrils and ribbons

Here, we have successfully produced ribbons and fibrils as described in the literature^15, 24^. Of note, we observed that different batches of assemblies obtained in the low salt conditions displayed different digestion patterns and only some of them corresponded to the reported ribbon fold (Supporting Figure 7). This was not the case for the fibrils which reproducibly gave the same PK digestion pattern and structural characteristics. This is consistent with recent studies that showed that different polymorphs can in some cases be generated stochastically in the same experimental conditions^38–40^. Some low salt polymorphs had a more fibril-like profile, based on PK digestion pattern, but remained less susceptible to undergo secondary nucleation compared to fibrils (see Supporting Figure 8).

The fact that we never observed a banding pattern corresponding to the fibrils upon PK digestion of the LSxH polymorphs (ribbons seeded in buffer promoting fibrils) was unexpected. Indeed, fibrils form faster than ribbons *de novo* and fibrils also amplify more efficiently than ribbons. We would therefore expect that primary nucleation in high salt would produce some fibrils that would quickly represent a sizable proportion of the final product (see below). Our findings indicate that the presence of seeds is very efficient at blocking primary nucleation given the ability of fibrils to become the dominant species upon amplification (Figure 5).

### Cellular effects of fibrils and ribbons

We tested the effects of the generated polymorphs on cells. As reported^24, 41^, fibrils strongly induce aggregation in treated cells compared to ribbons, however, we did not notice differences in toxicity between the polymorphs, based on measuring the levels of lactate dehydrogenase at days 4, 7 and 10. There was no significant difference in cytotoxicity between HS/LS treated cells and non-treated cells, indicating the HS/LS did not induce cytotoxicity on D4/7/10. Bousset et al.^24^ reported that fibrils were noticeably more toxic than ribbons in their assay, which examined cytotoxicity at early time points (24 h - 48 h treatment). In their case, there was a clear dose-dependent decrease in cell survival measured by cell counting and 3-(4,5-dimethylthiazol-2-yl)-2,5-diphenyltetrazolium bromide (MTT) assay, in undifferentiated SH-SY5Y cells. They further showed induction of apoptosis in differentiated cells, within 48 h. As our first time point is at day 4, it is possible that we did not observe this early apoptotic phase, which might be linked to the interaction between the αSyn aggregates and the membrane. Based on our data, it seems that once αSyn fibrils and ribbons have been taken up by the cells, they induce endogenous αSyn aggregation without a noticeable effect on toxicity^32^.

### Fibrils undergo secondary elongation more readily than ribbons

Our single molecule data revealed a striking difference in the mechanism of amyloid growth between fibrils and ribbons. Our hypothesis that the observed increase in the residence time was due to the formation of 3D-aggregates was validated using high-resolution microscopy, showing branched networks of filaments upon amplification of fibrils. The propensity of fibrils to grow by secondary nucleation was not limited to our set of experimental conditions. Rather, the same properties were seen using other sets of experimental conditions (monomer, buffer, temperature). The conditions of amplification could however strongly influence the mechanism of growth. In conditions known to promote secondary nucleation (e.g., low pH, high salt in condition B) both polymorphs amplified through 3D-growth. On the other hand, amplification at low concentration of monomers prevented secondary nucleation, as expected. Yet, in both cases, fibrils remained more efficient at amplifying than ribbons.

Buell et al.^4^ analyzed in detail how solution conditions affect the kinetics of these steps. They showed that in acidic quiescent conditions (pH < 6), seeded aggregation of αSyn was much faster than at physiological pH, due to a dramatic increase in the rate of secondary nucleation^35^. Peduzzo et al. ^42^ then studied the seeded amplification of ribbons and fibrils in acidic conditions (pH 6.5 or 5.5). They found that at low seed concentration (which favors secondary nucleation), fibrils and ribbons showed different amplification profiles, suggesting different relative contributions of elongation and secondary nucleation to the growth of αSyn assemblies. Amplified fibrils also looked more branched on atomic force spectroscopy images than amplified ribbons that remain mainly linear. The branching phenotype was lost in reaction where higher seed concentrations were used, which limits secondary nucleation. The authors concluded that ribbons and fibrils may be different “in the relative ability to act as templates for fibril elongation and secondary nucleation”. Our data support this conclusion and show that this difference in behavior holds true for a range of conditions.

### Aggregate conformation as a determinant of secondary nucleation

The nucleation and growth processes of αSyn aggregation have been well characterized. Different studies have established that secondary nucleation is the leading process of amplification for αSyn at pH < 6.0^4–35,42^ as well as for other amyloid-forming proteins (insulin, islet amyloid polypeptide (IAPP) or Amyloid β). Secondary nucleation is a multi-step process where monomers initially attach to the surface then convert into an aggregation-competent fold (nucleus formation) and elongate. Kumari et al. ^43^ showed that the positively charged N-terminus of αSyn monomers could interact with the negatively charged C-terminal domain of the aggregated αSyn. Removing the electrostatic charges by truncations of the C-terminus of αSyn allow secondary nucleation to become the main growth mechanism at neutral pH^44^. It has been reported that the physical size of the seed could influence its ability to undergo secondary nucleation^45^ but in our hands, sonicated ribbons were longer than the sonicated fibrils.

As a surface-catalyzed process, it is not surprising in retrospect that strains would be prone to different growth mechanisms. Cross-reactivity between Aβ40 and Aβ42 has been studied, and despite their high identity, Aβ40 and Aβ42 form strikingly different structures^46^. Separate studies have shown that Aβ40 fibrils could not only induce the formation of Aβ42 fibrils by surface-catalyzed secondary nucleation while Aβ42 fibrils did support this cross-nucleation^47, 48^. In the case of αSyn, polymorphic assemblies possess different core structures and expose different residues at their surface, evidenced by variation in conformational antibody recognition, for example^49^. NMR shows that the N-terminus of ribbons tends to be more structured than in fibrils^50^, making it less solvent-accessible and therefore resistant to proteolysis^49^. In these studies, the C-terminal domain of αSyn remained dynamic and unresolved, but it is likely that the structure of the amyloid core will have a long-range influence on the rest of the protein as well. A recent study by Pálmadottir et al.^51^ illustrates the influence of polymorphs on monomer-aggregate interactions. Polymorphs generated in the same conditions (10 mM Tris, pH 7.0) could adopt different morphologies and displayed different binding modes to αSyn monomers, as assessed by NMR. One polymorph bound to the N-terminus of the monomer as previously described^35,43^, while the other interacted with a large part of the monomer. A similar mechanism could be at play here.

### Impact of secondary nucleation on strain retention

With the development of αSyn SAA for the study of synucleinopathies^52^, the question of whether structural characteristics of the seeds are retained upon amplification becomes very relevant. More studies are still needed to get a definitive answer, but it is thought for now that the assemblies generated by secondary nucleation are more influenced by the reaction conditions than by the parent seed. This was shown for αSyn, insulin or the mouse prion protein^48^. The case of Aβ is less clear, as the assemblies’ surface selectively interacts with some isomers that are structurally compatible^53^, yet the secondary nuclei do not necessarily inherit the properties of the seed^47^. Here, we used our model to ask whether the difference of growth mechanism could have an impact on the final product of the amplification, *irrespective of strain retention*. Indeed, this is relevant if we are to use *in vitro* amplification to study the heterogeneity of αSyn strains in patients. To do this, we simulated a competition between different strains, with two different dynamics of growth (as calculated in Figure 5B). The initial conditions were varied by changing the ratio of ribbons to fibrils and growth was simulated for up to 80 h (Figure 5D). Adding as little as 10 parts per billion (0.0000001%) of fibrils would already have a significant impact on the final composition of the solution (55.9% of ribbons final). Adding > 0.1% of fibrils lead to final solutions where the presence of ribbons becomes almost insignificant (< 5%). Obviously, this simulation did not account for many other variables and processes that are at play during *in vitro* amplification and is based on overly contrasted differences of mechanisms between the strains. However, when we performed RT-QuIC assays with ribbons and fibrils seeds at the same time, minute amounts of fibrils dramatically changed the response of the assay (Figure 5F). This suggests that differences in mechanisms of growth between strains should be kept in mind for future studies of pathological αSyn aggregates relying on amplification.

In conclusion, we demonstrated that αSyn polymorphism could impact the mode of amyloid amplification. The ability of fibrils, but not ribbons, to support secondary nucleation at neutral pH leads to faster amplification *in vitro* as well as the formation of larger aggregates in cells. This should be considered when comparing different αSyn polymorphs and strains linked to different pathologies that would presumably have notable disparities in properties and conformations.

## References

1. Goedert, M.; Jakes, R.; Spillantini, M. G., The Synucleinopathies: Twenty Years On. J Parkinsons Dis 2017, 7 (s1), S51–S69.

2. Ma, J.; Gao, J.; Wang, J.; Xie, A., Prion-Like Mechanisms in Parkinson’s Disease. Frontiers in neurology 2019, 13 (552).

3. Ke, P. C.; Zhou, R.; Serpell, L. C.; Riek, R.; Knowles, T. P. J.; Lashuel, H. A.; Gazit, E.; Hamley, I. W.; Davis, T. P.; Fändrich, M.; Otzen, D. E.; Chapman, M. R.; Dobson, C. M.; Eisenberg, D. S.; Mezzenga, R., Half a century of amyloids: past, present and future. Chemical Society reviews 2020, 49 (15), 5473–5509.

4. Buell, A. K.; Galvagnion, C.; Gaspar, R.; Sparr, E.; Vendruscolo, M.; Knowles, T. P.; Linse, S.; Dobson, C. M., Solution conditions determine the relative importance of nucleation and growth processes in α-synuclein aggregation. Proceedings of the National Academy of Sciences of the United States of America 2014, 111 (21), 7671–6.

5. Russo, M. J.; Orru, C. D.; Concha-Marambio, L.; Giaisi, S.; Groveman, B. R.; Farris, C. M.; Holguin, B.; Hughson, A. G.; LaFontant, D. E.; Caspell-Garcia, C.; Coffey, C. S.; Mollon, J.; Hutten, S. J.; Merchant, K.; Heym, R. G.; Soto, C.; Caughey, B.; Kang, U. J., High diagnostic performance of independent alpha-synuclein seed amplification assays for detection of early Parkinson’s disease. Acta neuropathologica communications 2021, 9 (1), 179.

6. Ferreira, N. D. C.; Caughey, B., Proteopathic Seed Amplification Assays for Neurodegenerative Disorders. Clinics in laboratory medicine 2020, 40 (3), 257–270.

7. Siderowf, A.; Concha-Marambio, L.; Lafontant, D. E.; Farris, C. M.; Ma, Y.; Urenia, P. A.; Nguyen, H.; Alcalay, R. N.; Chahine, L. M.; Foroud, T.; Galasko, D.; Kieburtz, K.; Merchant, K.; Mollenhauer, B.; Poston, K. L.; Seibyl, J.; Simuni, T.; Tanner, C. M.; Weintraub, D.; Videnovic, A.; Choi, S. H.; Kurth, R.; Caspell-Garcia, C.; Coffey, C. S.; Frasier, M.; Oliveira, L. M. A.; Hutten, S. J.; Sherer, T.; Marek, K.; Soto, C., Assessment of heterogeneity among participants in the Parkinson’s Progression Markers Initiative cohort using α-synuclein seed amplification: a cross-sectional study. The Lancet. Neurology 2023, 22 (5), 407–417.

8. Groveman, B. R.; Orrù, C. D.; Hughson, A. G.; Raymond, L. D.; Zanusso, G.; Ghetti, B.; Campbell, K. J.; Safar, J.; Galasko, D.; Caughey, B., Rapid and ultra-sensitive quantitation of disease-associated α-synuclein seeds in brain and cerebrospinal fluid by αSyn RT-QuIC. Acta neuropathologica communications 2018, 6 (1), 7.

9. Iranzo, A.; Fairfoul, G.; Ayudhaya, A. C. N.; Serradell, M.; Gelpi, E.; Vilaseca, I.; Sanchez-Valle, R.; Gaig, C.; Santamaria, J.; Tolosa, E.; Riha, R. L.; Green, A. J. E., Detection of α-synuclein in CSF by RT-QuIC in patients with isolated rapid-eye-movement sleep behaviour disorder: a longitudinal observational study. The Lancet. Neurology 2021, 20 (3), 203–212.

10. Miglis, M. G.; Adler, C. H.; Antelmi, E.; Arnaldi, D.; Baldelli, L.; Boeve, B. F.; Cesari, M.; Dall’Antonia, I.; Diederich, N. J.; Doppler, K.; Dušek, P.; Ferri, R.; Gagnon, J. F.; Gan-Or, Z.; Hermann, W.; Högl, B.; Hu, M. T.; Iranzo, A.; Janzen, A.; Kuzkina, A.; Lee, J. Y.; Leenders, K. L.; Lewis, S. J. G.; Liguori, C.; Liu, J.; Lo, C.; Ehgoetz Martens, K. A.; Nepozitek, J.; Plazzi, G.; Provini, F.; Puligheddu, M.; Rolinski, M.; Rusz, J.; Stefani, A.; Summers, R. L. S.; Yoo, D.; Zitser, J.; Oertel, W. H., Biomarkers of conversion to α-synucleinopathy in isolated rapid-eye-movement sleep behaviour disorder. The Lancet. Neurology 2021, 20 (8), 671–684.

11. Rossi, M.; Baiardi, S.; Teunissen, C. E.; Quadalti, C.; van de Beek, M.; Mammana, A.; Maserati, M. S.; Van der Flier, W. M.; Sambati, L.; Zenesini, C.; Caughey, B.; Capellari, S.; Lemstra, A.; Parchi, P., Diagnostic Value of the CSF α-Synuclein Real-Time Quaking-Induced Conversion Assay at the Prodromal MCI Stage of Dementia With Lewy Bodies. Neurology 2021.

12. Wood, H., A seed amplification assay to detect Parkinson disease pathology. Nature reviews. Neurology 2023.

13. Berg, D.; Klein, C., α-synuclein seed amplification and its uses in Parkinson’s disease. The Lancet. Neurology 2023, 22 (5), 369–371.

14. Ayers, J. I.; Lee, J.; Monteiro, O.; Woerman, A. L.; Lazar, A. A.; Condello, C.; Paras, N. A.; Prusiner, S. B., Different α-synuclein prion strains cause dementia with Lewy bodies and multiple system atrophy. Proceedings of the National Academy of Sciences of the United States of America 2022, 119 (6).

15. Van der Perren, A.; Gelders, G.; Fenyi, A.; Bousset, L.; Brito, F.; Peelaerts, W.; Van den Haute, C.; Gentleman, S.; Melki, R.; Baekelandt, V., The structural differences between patient-derived α-synuclein strains dictate characteristics of Parkinson’s disease, multiple system atrophy and dementia with Lewy bodies. Acta Neuropathol 2020, 139 (6), 977–1000.

16. Holec, S. A. M.; Woerman, A. L., Evidence of distinct α-synuclein strains underlying disease heterogeneity. Acta Neuropathol 2021, 142 (1), 73–86.

17. Carta, M.; Aguzzi, A., Molecular foundations of prion strain diversity. Current opinion in neurobiology 2021, 72, 22–31.

18. Lau, H. H. C.; Ingelsson, M.; Watts, J. C., The existence of Aβ strains and their potential for driving phenotypic heterogeneity in Alzheimer’s disease. Acta Neuropathologica 2020.

19. Narasimhan, S.; Guo, J. L.; Changolkar, L.; Stieber, A.; McBride, J. D.; Silva, L. V.; He, Z.; Zhang, B.; Gathagan, R. J.; Trojanowski, J. Q.; Lee, V. M. Y., Pathological Tau Strains from Human Brains Recapitulate the Diversity of Tauopathies in Nontransgenic Mouse Brain. The Journal of neuroscience : the official journal of the Society for Neuroscience 2017, 37 (47), 11406–11423.

20. Porta, S.; Xu, Y.; Lehr, T.; Zhang, B.; Meymand, E.; Olufemi, M.; Stieber, A.; Lee, E. B.; Trojanowski, J. Q.; Lee, V. M., Distinct brain-derived TDP-43 strains from FTLD-TDP subtypes induce diverse morphological TDP-43 aggregates and spreading patterns in vitro and in vivo. Neuropathology and applied neurobiology 2021, 47 (7), 1033–1049.

21. Yang, Y.; Shi, Y.; Schweighauser, M.; Zhang, X.; Kotecha, A.; Murzin, A. G.; Garringer, H. J.; Cullinane, P. W.; Saito, Y.; Foroud, T.; Warner, T. T.; Hasegawa, K.; Vidal, R.; Murayama, S.; Revesz, T.; Ghetti, B.; Hasegawa, M.; Lashley, T.; Scheres, S. H. W.; Goedert, M., Structures of α-synuclein filaments from human brains with Lewy pathology. Nature 2022, 610 (7933), 791–795.

22. Shahnawaz, M.; Mukherjee, A.; Pritzkow, S.; Mendez, N.; Rabadia, P.; Liu, X.; Hu, B.; Schmeichel, A.; Singer, W.; Wu, G.; Tsai, A. L.; Shirani, H.; Nilsson, K. P. R.; Low, P. A.; Soto, C., Discriminating α-synuclein strains in Parkinson’s disease and multiple system atrophy. Nature 2020, 578 (7794), 273–277.

23. Schweighauser, M.; Shi, Y.; Tarutani, A.; Kametani, F.; Murzin, A. G.; Ghetti, B.; Matsubara, T.; Tomita, T.; Ando, T.; Hasegawa, K.; Murayama, S.; Yoshida, M.; Hasegawa, M.; Scheres, S. H. W.; Goedert, M., Structures of α-synuclein filaments from multiple system atrophy. Nature 2020, 585 (7825), 464–469.

24. Bousset, L.; Pieri, L.; Ruiz-Arlandis, G.; Gath, J.; Jensen, P. H.; Habenstein, B.; Madiona, K.; Olieric, V.; Böckmann, A.; Meier, B. H.; Melki, R., Structural and functional characterization of two alpha-synuclein strains. Nature communications 2013, 4, 2575.

25. Peelaerts, W.; Bousset, L.; Van der Perren, A.; Moskalyuk, A.; Pulizzi, R.; Giugliano, M.; Van den Haute, C.; Melki, R.; Baekelandt, V., α-Synuclein strains cause distinct synucleinopathies after local and systemic administration. Nature 2015, 522 (7556), 340–4.

26. Suzuki, G.; Imura, S.; Hosokawa, M.; Katsumata, R.; Nonaka, T.; Hisanaga, S. I.; Saeki, Y.; Hasegawa, M., α-synuclein strains that cause distinct pathologies differentially inhibit proteasome. eLife 2020, 9.

27. Torre-Muruzabal, T.; Van der Perren, A.; Coens, A.; Gelders, G.; Barber Janer, A.; Camacho-Garcia, S.; Klingstedt, T.; Nilsson, P.; Stefanova, N.; Melki, R.; Baekelandt, V.; Peelaerts, W., Host oligodendrogliopathy and ɑ-synuclein strains dictate disease severity in multiple system atrophy. Brain : a journal of neurology 2022.

28. Fayard, A.; Fenyi, A.; Lavisse, S.; Dovero, S.; Bousset, L.; Bellande, T.; Lecourtois, S.; Jouy, C.; Guillermier, M.; Jan, C.; Gipchtein, P.; Dehay, B.; Bezard, E.; Melki, R.; Hantraye, P.; Aron Badin, R., Functional and neuropathological changes induced by injection of distinct alpha-synuclein strains: A pilot study in non-human primates. Neurobiology of disease 2023, 180, 106086.

29. Bhumkar, A.; Magnan, C.; Lau, D.; Jun, E. S. W.; Dzamko, N.; Gambin, Y.; Sierecki, E., Single-Molecule Counting Coupled to Rapid Amplification Enables Detection of α-Synuclein Aggregates in Cerebrospinal Fluid of Parkinson’s Disease Patients. Angewandte Chemie (International ed. in English) 2021, 60 (21), 11874–11883.

30. Lau, D.; Magnan, C.; Hill, K.; Cooper, A.; Gambin, Y.; Sierecki, E., Single Molecule Fingerprinting Reveals Different Amplification Properties of α-Synuclein Oligomers and Preformed Fibrils in Seeding Assay. ACS chemical neuroscience 2022, 13 (7), 883–896.

31. LeVine, H., 3rd, Thioflavine T interaction with synthetic Alzheimer’s disease beta-amyloid peptides: detection of amyloid aggregation in solution. Protein science : a publication of the Protein Society 1993, 2 (3), 404–10.

32. Gao, J.; Perera, G.; Bhadbhade, M.; Halliday, G. M.; Dzamko, N., Autophagy activation promotes clearance of α-synuclein inclusions in fibril-seeded human neural cells. The Journal of biological chemistry 2019, 294 (39), 14241–14256.

33. Brown, J. W. P.; Bauer, A.; Polinkovsky, M. E.; Bhumkar, A.; Hunter, D. J. B.; Gaus, K.; Sierecki, E.; Gambin, Y., Single-molecule detection on a portable 3D-printed microscope. Nature communications 2019, 10 (1), 5662.

34. Knowles, T. P.; Waudby, C. A.; Devlin, G. L.; Cohen, S. I.; Aguzzi, A.; Vendruscolo, M.; Terentjev, E. M.; Welland, M. E.; Dobson, C. M., An analytical solution to the kinetics of breakable filament assembly. *Science (New York*, N.Y.) 2009, 326 (5959), 1533–7.

35. Gaspar, R.; Meisl, G.; Buell, A. K.; Young, L.; Kaminski, C. F.; Knowles, T. P. J.; Sparr, E.; Linse, S., Secondary nucleation of monomers on fibril surface dominates α-synuclein aggregation and provides autocatalytic amyloid amplification. Quarterly reviews of biophysics 2017, 50, e6.

36. Ramis, R.; Ortega-Castro, J.; Vilanova, B.; Adrover, M.; Frau, J., Unraveling the NaCl Concentration Effect on the First Stages of α-Synuclein Aggregation. Biomacromolecules 2020, 21 (12), 5200–5212.

37. Gaspar, R.; Lund, M.; Sparr, E.; Linse, S., Anomalous Salt Dependence Reveals an Interplay of Attractive and Repulsive Electrostatic Interactions in α-synuclein Fibril Formation. QRB discovery 2020, 1, e2.

38. De Giorgi, F.; Laferrière, F.; Zinghirino, F.; Faggiani, E.; Lends, A.; Bertoni, M.; Yu, X.; Grélard, A.; Morvan, E.; Habenstein, B.; Dutheil, N.; Doudnikoff, E.; Daniel, J.; Claverol, S.; Qin, C.; Loquet, A.; Bezard, E.; Ichas, F., Novel self-replicating α-synuclein polymorphs that escape ThT monitoring can spontaneously emerge and acutely spread in neurons. 2020, 6 (40), eabc4364.

39. Pálmadóttir, T.; Waudby, C. A.; Bernfur, K.; Christodoulou, J.; Linse, S.; Malmendal, A., Morphology-Dependent Interactions between α-Synuclein Monomers and Fibrils. International journal of molecular sciences 2023, 24 (6).

40. Singh, B. P.; Morris, R. J.; Kunath, T.; MacPhee, C. E.; Horrocks, M. H., Lipid-induced polymorphic amyloid fibril formation by α-synuclein. Protein science : a publication of the Protein Society 2023, e4736.

41. Tittelmeier, J.; Druffel-Augustin, S.; Alik, A.; Melki, R.; Nussbaum-Krammer, C., Dissecting aggregation and seeding dynamics of α-Syn polymorphs using the phasor approach to FLIM. Commun Biol 2022, 5 (1), 1345.

42. Peduzzo, A.; Linse, S.; Buell, A. K., The Properties of α-Synuclein Secondary Nuclei Are Dominated by the Solution Conditions Rather than the Seed Fibril Strain. ACS chemical neuroscience 2020, 11 (6), 909–918.

43. Kumari, P.; Ghosh, D.; Vanas, A.; Fleischmann, Y.; Wiegand, T.; Jeschke, G.; Riek, R.; Eichmann, C., Structural insights into α-synuclein monomer-fibril interactions. Proceedings of the National Academy of Sciences of the United States of America 2021, 118 (10).

44. van der Wateren, I. M.; Knowles, T. P. J.; Buell, A. K.; Dobson, C. M.; Galvagnion, C., C-terminal truncation of α-synuclein promotes amyloid fibril amplification at physiological pH. Chemical science 2018, 9 (25), 5506–5516.

45. Sakunthala, A.; Datta, D.; Navalkar, A.; Gadhe, L.; Kadu, P.; Patel, K.; Mehra, S.; Kumar, R.; Chatterjee, D.; Devi, J.; Sengupta, K.; Padinhateeri, R.; Maji, S. K., Direct Demonstration of Seed Size-Dependent α-Synuclein Amyloid Amplification. The journal of physical chemistry letters 2022, 13 (28), 6427–6438.

46. Cukalevski, R.; Yang, X.; Meisl, G.; Weininger, U.; Bernfur, K.; Frohm, B.; Knowles, T. P. J.; Linse, S., The Aβ40 and Aβ42 peptides self-assemble into separate homomolecular fibrils in binary mixtures but cross-react during primary nucleation. Chemical science 2015, 6 (7), 4215–4233.

47. Brännström, K.; Islam, T.; Gharibyan, A. L.; Iakovleva, I.; Nilsson, L.; Lee, C. C.; Sandblad, L.; Pamrén, A.; Olofsson, A., The Properties of Amyloid-β Fibrils Are Determined by their Path of Formation. Journal of molecular biology 2018, 430 (13), 1940–1949.

48. Hadi Alijanvand, S.; Peduzzo, A.; Buell, A. K., Secondary Nucleation and the Conservation of Structural Characteristics of Amyloid Fibril Strains. Frontiers in molecular biosciences 2021, 8, 669994.

49. Landureau, M.; Redeker, V.; Bellande, T.; Eyquem, S.; Melki, R., The differential solvent exposure of N-terminal residues provides “fingerprints” of alpha-synuclein fibrillar polymorphs. J Biol Chem 2021, 296, 100737.

50. Gath, J.; Bousset, L.; Habenstein, B.; Melki, R.; Böckmann, A.; Meier, B. H., Unlike twins: an NMR comparison of two α-synuclein polymorphs featuring different toxicity. PloS one 2014, 9 (3), e90659.

51. Pálmadóttir, T.; Waudby, C. A.; Bernfur, K.; Christodoulou, J.; Linse, S.; Malmendal, A., Morphology-Dependent Interactions between alpha-Synuclein Monomers and Fibrils. 2023, 24 (6), 5191.

52. Standke, H. G.; Kraus, A., Seed amplification and RT-QuIC assays to investigate protein seed structures and strains. Cell and tissue research 2023, 392 (1), 323–335.

53. Thacker, D.; Sanagavarapu, K.; Frohm, B.; Meisl, G.; Knowles, T. P. J.; Linse, S., The role of fibril structure and surface hydrophobicity in secondary nucleation of amyloid fibrils. Proceedings of the National Academy of Sciences of the United States of America 2020, 117 (41), 25272–25283.

