## Supplemental Information for "Single molecule fingerprinting reveals different growth mechanisms in seed amplification assays for different polymorphs of αSynuclein fibrils"

### MATERIALS AND METHODS

#### Expression and purification of human α-syn wild-type and K23Q.

The expression and purification of untagged recombinant human α-syn wild-type (WT) was performed as described in Lau et al.[1] Purified α-syn were checked by reducing-SDS-PAGE and had its concentration determined spectroscopically at 280 nm absorbance with ε = 5960 $M^{-1}cm^{-1}$. The expression and purification of recombinant human α-syn K23Q was performed as described in Groveman et al 2018 Purified proteins were aliquoted, flash frozen with liquid nitrogen for storage at -80 °C.

#### Production recombinant α-syn aggregates.

Purified recombinant human α-syn WT was exchanged into a low salt buffer (LSB, 5 mM Tris, pH 7.5, 0.02% w/v $NaN_3$) using Zeba spin columns (Thermo Scientific, 89890).
Production of recombinant low salt and high salt α-syn aggregates were performed in screw top microtubes (SSIbio, tubes 2340-00 and caps 2001-00) with a final volume of 600 µL. Generation of low salt ribbons was initiated by diluting human α-syn wild-type (250 µM) in the low salt buffer (5 mM Tris, pH 7.5, 0.02% w/v $NaN_3$). Generation of high salt fibrils was initiated by diluting human α-syn wild-type (250 µM) in the high salt buffer (HSB, 50 mM Tris, pH 7.5, 150 mM KCl, 0.02% w/v $NaN_3$). The tubes were shaken horizontally at 600 rpm, 37 °C inside a falcon tube strapped on a vortexer. Ribbons and fibrils α-syn assemblies were collected after 7 and 2 days respectively. Ribbons and fibrils were sonicated on the M220 focused-ultrasonicator (Covaris) with a set point of 12 °C with peak power of 75 watts, duty factor of 25, 200 cycles/burst, delay 5 sec, 60 cycles. Non-sonicated and sonicated materials were frozen in liquid nitrogen for storage at -80 °C. Samples were thawed on tap water and kept at room temperature to prevent disassembly at 4°C.

#### Cross-seeding α-syn aggregates in low salt and high salt buffers.

Ribbons and fibrils aggregates (250 µM) were sonicated for 5 min at room temperature on a Branson sonicator with setting 3, 30% duty cycle. Seeding reaction were initiated by adding 5 µL of the sonicated materials to low salt buffer (LSB, 5 mM Tris, pH 7.5, 0.02% w/v $NaN_3$) or high salt buffer (HSB, 50 mM Tris, pH 7.5, 150 mM KCl, 0.02% w/v $NaN_3$) containing human α-syn wild-type (100 µM) with a final volume of 100 µL as the first batch. Assembly was induced by shaking at 180 rpm, 37 °C for 8 days. Ribbons seeded in LSB (LSxL), ribbons seeded in HSB (LSxH), fibrils seeded in LSB (HSxL) and fibrils seeded in HSB (HSxH) were sonicated on the M220 focused-ultrasonicator (Covaris) with a set point of 12 °C with peak power of 75 watts, duty factor of 25, 200 cycles/burst, delay 5 sec, 60 cycles. The second batch of cross-seeded aggregates used for cell studies were re-prepared at final volume of 500 µL and sonicated (Qsonica, Q500) at 50% strength, 1 min sonication time with 3 s on/off. Non-sonicated and sonicated materials were frozen in liquid nitrogen for storage at -80 °C. Samples were thawed on tap water and kept at room temperature to prevent disassembly at 4 °C. Two batches of ribbon seeded in LSB (LSxL1 and LSxL2) were produced.

#### Negative staining electron microscopy.

α-syn ribbons, fibrils and cross-seeded α-syn aggregates were diluted to 12.5 µM with sterile MilliQ water. Samples were negatively stained by adapting a published protocol.[1] A copper grid (200 mesh, coated with carbon and Formvar, Ted Pella, 01811) was cleaned by glow discharge and the sample was applied onto with 1 min incubation at room temperature. The grid was wicked dry before staining uranyl acetate (2% w/v) for 30 sec. This process was repeated two more times and the grid was air dried. Micrographs were collected using a FEI

Tecnai G2 20 electron microscope at 19500-fold magnification mounted with a Gatan Orius SC1000A CCD camera. Particle diameters were measured using ImageJ.

**Proteinase K digestion and reducing SDS-PAGE.**

Non-sonicated ribbons and fibrils (40 µL of 250 µM) were centrifuged at 50000 g for 30 min at 25 °C (S100AT3, Eppendorf Himac). The concentration of α-syn in the supernatant was determined spectroscopically at 280 nm to estimate the concentration of α-syn in the pellet. Ribbons and fibrils were diluted to 40 µM (monomer equivalent) in the digestion buffer to a final volume of 10 µL. Cross-seeded α-syn aggregates (100 µM) had the assemblies diluted to a final concentration of 40 µM without centrifugation to a final volume of 10 µL for digestion. Monomeric α-syn WT (40 µM) were used as control to ensure the proteinase K was functional. For each reaction, 5 µL was withdrawn and quenched in a Tricine SDS sample buffer (Thermo Scientific, LC1676) containing β-mercaptoethanol (4% v/v). The remaining solution was digested by adding 0.21 µL of 37.5 µg/mL of proteinase K (final 1.5 µg/mL), incubated at 37 °C for exactly 5, 15 or 30 min at 37 °C and quenched by adding an equal volume of Tricine running buffer containing β-mercaptoethanol (4% v/v). All samples were then heated at 90 °C for 10 min and resolved using reducing SDS-PAGE with Tricine gels (Thermo Scientific, EC66955BOX) at 120 V for 1 h 50 min. Gels were fixed with a fixing solution (50% v/v methanol, 7% v/v acetic acid) for 15 min, stained with SYPRO Ruby stain (Thermo Scientific, S12000) overnight at 4 °C in the dark. Gel was destained with a wash-solution (10% v/v methanol, 7% v/v acetic acid) for 15 min and washed twice with MilliQ water before imaging using a GelDoc (Bio-Rad). Gel were analysed using ImageJ. Three independent repeats were performed for time course proteinase K digestion of ribbons and fibrils. Two independent repeats were performed for the proteolytic digestion of cross-seeded aggregates.

**Single molecule seed amplification assay measurements and analysis.**

Single molecule seed amplification assay (smSAA) of α-syn aggregates followed a previously established protocol.[2] Human α-syn K23Q were filtered using an 100k MWCO Amicon filter (Merck Millipore, UFC510096) to remove aggregates. Concentrations were determined spectroscopically with absorbance at 280 nm and ε = 5960 $M^{-1}cm^{-1}$. Sonicated ribbons (6.3 nM) and fibrils (1 nM) were diluted in phosphate buffer saline (PBS, 137 mM NaCl, 2.7 mM KCl, 10 mM $Na_2HPO_4$, and 1.8 mM $KH_2PO_4$), filtered human α-syn K23Q (20 µM) and thioflavin T (10 µM) to a final volume of 40 µL. This volume was separated into two equal volume reactions. One volume was amplified at 55 °C for 5 h in a thermocycler and the other volume was measured on the 3D confocal microscope immediately as control (before amplification). Samples were loaded to a custom polydimethylsiloxane (PDMS) plate adhered to a glass coverslip (ProSciTech, G425-4860) and observed using the inverted 3D printed confocal microscope, equipped with a 450 nm laser and water immersion 40×/1.2 NA objective (Zeiss).[2] Emitted fluorescence from ThT was filtered by a dichroic mirror (488 nm) and a long-pass filter (500 nm) before focusing onto a single photon avalanche diode (Micro Photon Devices). Fluorescence spectroscopy traces were recorded for 10 min, 100 s/trace in 10 ms bins. Traces acquired on different days were pooled together for analysis. Traces were analyzed using a custom python script on Spyder version 4.0.1 to extract single molecule parameters in an automated and unbiased manner as previously described.[1] SmSAA of sonicated ribbons (25 nM) and sonicated fibrils (1 nM) using concentration of human α-syn WT (0.88, 1.75, 3.5, 7, 14, 28 µM) were performed in PBS at 55 °C for 5 h.

Sonicated ribbons seeded in LSB (LSxL), ribbons seeded in HSB (LSxH), fibrils seeded in LSB (HSxL), fibrils seeded in HSB (HSxH) were diluted to 20, 20, 5 and 0.5 nM respectively in PBS containing filtered human α-syn K23Q (20 µM) and thioflavin T (10 µM) for smSAA. Reactions were amplified at 55 °C for 5 h. Sonicated cross-seeded aggregates used for cell-studies were amplified using the same protocol with an adjustment of final concentration of aggregates of 20, 10, 2 and 2 nM for LSxL, LSxH, HSxL, HSxH respectively. Traces were pooled together for analysis from at least two independent amplification experiments.

**Confocal microscopy on amplified α-syn fibrils.**

High salt fibrils were amplified using smSAA with PBS containing ThT (10 µM) and human α-syn K23Q at 55 °C for 3 h. Two reactions were set up. A small volume of amplified reaction (5 µL) was transferred between two clean coverslips and mounted onto a Chamlide chamber (Live Cell Instrument, CMB) for imaging using confocal microscopy. Confocal microscopy was performed on an LSM880 laser scanning confocal microscope with AiryScan (Zeiss, Oberkochen, Germany) using a 63×/1.4 oil immersion objective. ThT dye bound onto the amplified fibrils was excited with a 488 nm Argon laser using powers setting 20 and 30%. Fluorescence emission was cleaned up through a dual band emission filter (420-475 / 500-545 nm) and detected by an Airyscan GaAsP photomultiplier tube array at a voxel-size of 0.04×0.04×0.17 $\mu m^3$. The z-stacks of sizes between 8 and 10 µm were recorded to capture the whole structure of the fibrillar aggregate. Raw images stacks were AiryScan processed using Zen Black 2.3 (3D processing, auto filter). Movies and images were rendered using Imaris Microscopy Image Analysis Software (Oxford Instruments) and length and width of each aggregates were measured by ImageJ.

**SH-SY5Y cell culture and treatment with recombinant α-syn aggregates.**

Protein concentrations of α-syn aggregates were re-measured by Pierce bicinchoninic acid assay (Thermo Fisher Scientific, 23227) with BSA as standard to normalize the treatment to differentiated SH-SY5Y cells. SH-SY5Y human neuroblastoma cells were cultured in DMEM/Ham's F-12 (Thermo Fisher Scientific, 11320033) supplemented with 10% w/v low endotoxin fetal bovine serum (Thermo Fisher Scientific, 16000069), 2 mM L-glutamine (Thermo Fisher, 25030081) and 1% w/v penicillin-streptomycin (Thermo Fisher Scientific, 10378016). Cells were seeded on coverslips in 12-well plates ($2.5 \times 10^4$ cells/well) and incubated overnight at 37°C, 5% v/v $CO_2$ and 95% humidity. Differentiation was induced in DMEM/Ham's F-12 containing 1% fetal bovine serum, 1% w/v penicillin-streptomycin and 10 µM of retinoic acid (Sigma Alrich, R2625-100MG) for 7 days and was verified using brightfield microscopy. Immediately before treating the cells, α-syn aggregates was diluted in sterile Dulbecco's PBS (DPBS, Thermo Fisher Scientific, 14040133) to 0.1 mg/ml (6.9 µM), sonicated for 30 s at 40% amplitude and 1 s on/off pulse (Qsonica, Q125). Cells were then treated with 1 mL of different strains of α-syn at 5 µg/mL (0.35 µM) and washed with DPBS after 2 days. Cells were further maintained for up to 10 days with media changed every 3 days.

**Confocal fluorescence microscopy of SH-SY5Y cells treated with recombinant α-syn aggregates.**

Cells on coverslips were washed with PBS (Thermo Fisher Scientific, 18912014) and fixed with 4% w/v paraformaldehyde (Sigma Alrich, 158127-3KG) for 20 min. After washing again with PBS, cells were permeabilized with 0.3% v/v triton (Sigma Alrich, X100-500ML) for 15 min. Blocking was performed with 3% w/v BSA (Sigma Alrich, A9647-1KG) and 0.1% v/v Triton for 1 h. The primary antibody used was mouse monoclonal anti-α-syn (BD Biosciences, 610787; 1:400 dilution) diluted in 3% w/v BSA and 0.1% v/v triton and incubated overnight at 4 °C. After washing with PBS, donkey anti-mouse Alexa Fluor 488 secondary antibodies (Invitrogen, A21202; 1:400 dilution) were added for 1 h in the dark. Following another wash with PBS, DAPI (Sigma Alrich, D9542-5MG) was added to the last wash at a 1:10,000 dilution and incubated for 15 min. Finally, coverslips were mounted with fluorescent mounting medium (Dako, S3023) and allowed to dry for 1 day before imaging using a Nikon C2 confocal microscope taken at 40× magnification. Images were analyzed for α-synuclein inclusions using ImageJ. The threshold tool was used to highlight stained areas and measure integrated density. Cell number was obtained based on DAPI staining. Cells close to the field of view (FOV) edges were excluded for analysis. Five images (30-100 cells per image) were analyzed per treatment condition. Fluorescent intensity was normalized to DAPI count for the number of SH-SY5Y cells. Statistical analysis was conducted using Prism software (GraphPad Software). Comparisons between two groups were performed using Student's t-test, while one-way ANOVA with Tukey's post-hoc test was used for comparisons among multiple groups. A p-value $< 0.05$ was considered statistically significant, and all figures show mean ± standard deviation (SD) of the data.

**Supporting Figure 1**

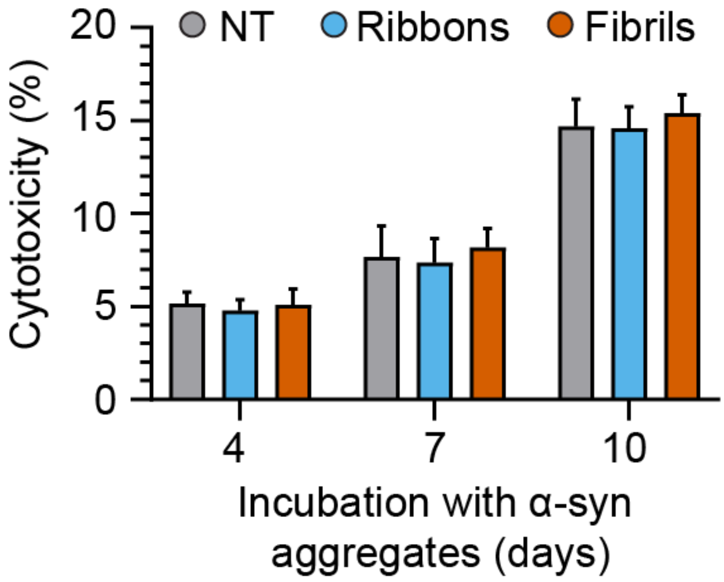


**Supporting Figure 1. Cytotoxicity in SH-SY5Y cells after incubation with ribbons or fibrils.**

Lactate dehydrogenase levels in SH-SY5Y cells were determined using the colorimetric cytotoxicity assay after treatment with 5 μg/mL ribbons or fibrils α-syn. Cytotoxicity was assessed at 4, 7 and 10 days after treatment. Bar graph of average percentage of cytotoxicity to maximum lactate dehydrogenase release. Error bars represent mean ± standard deviation.

**Supporting Figure 2**

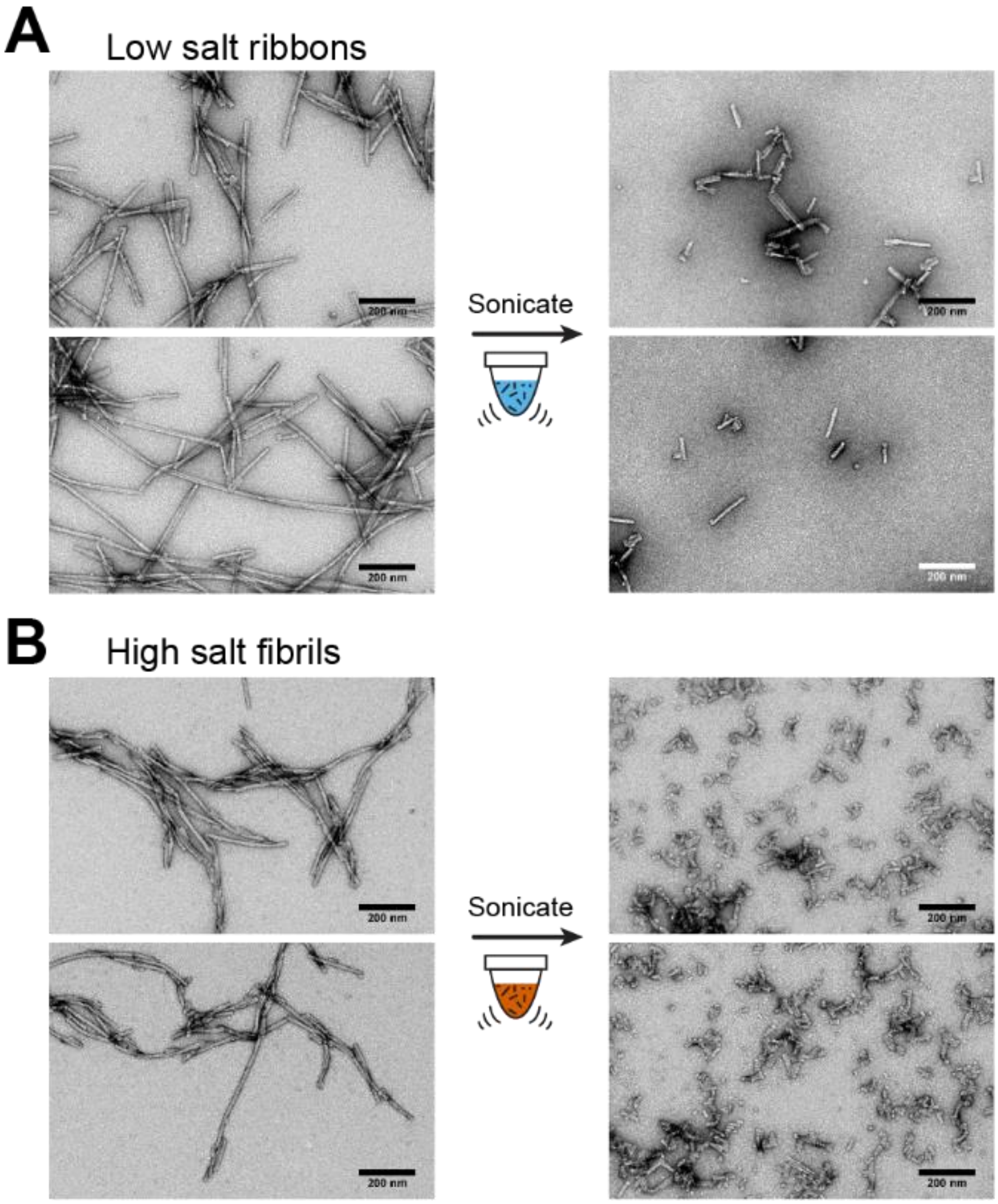


**Supporting Figure 2. Effect of sonication of recombinant α-syn aggregates for single molecule profiling.**

**A.** Electron micrographs of low salt ribbons before (left) and after (right) sonication with the M220 Covaris ultrasonicator. **B.** Electron micrographs of high salt fibrils before (left) and after (right) sonication. Scale bar 200 nm.

**Supporting Figure 3**

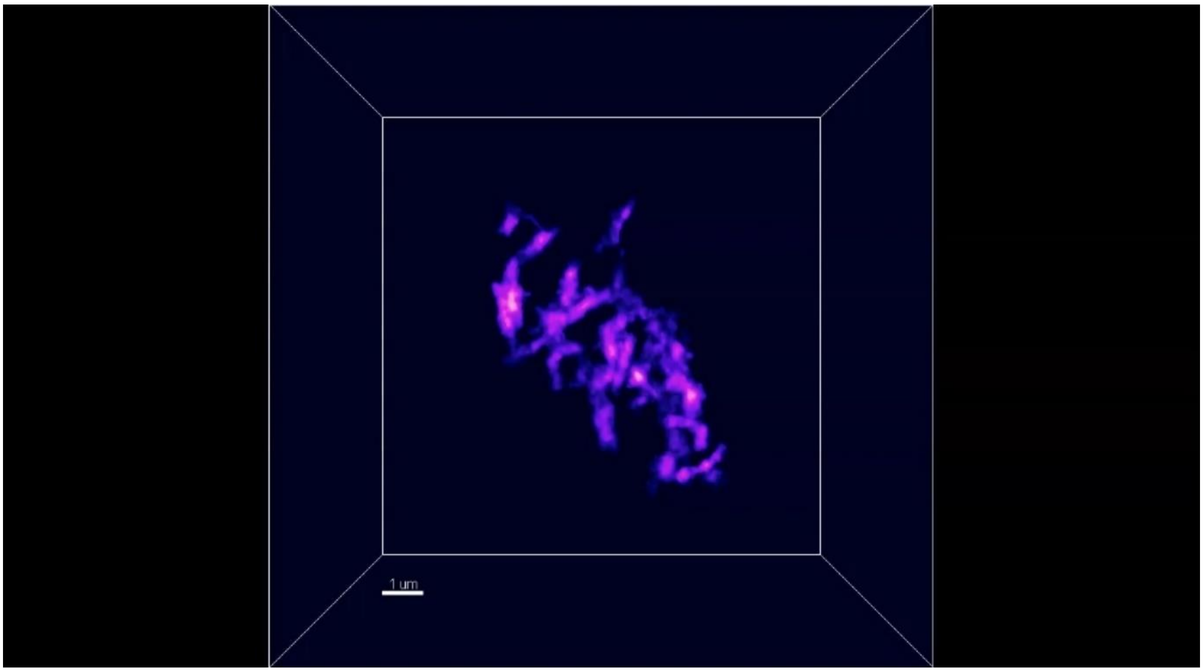


**Supporting Figure 3. 3D movie render of amplified high salt fibrils.**

Confocal microscopy images were collated to produce a movie. The aggregate was observed in reaction amplified at 55°C for 5 h in PBS with α-syn K23Q from fibrils as shown in Figure 4.

**Supporting Figure 4**

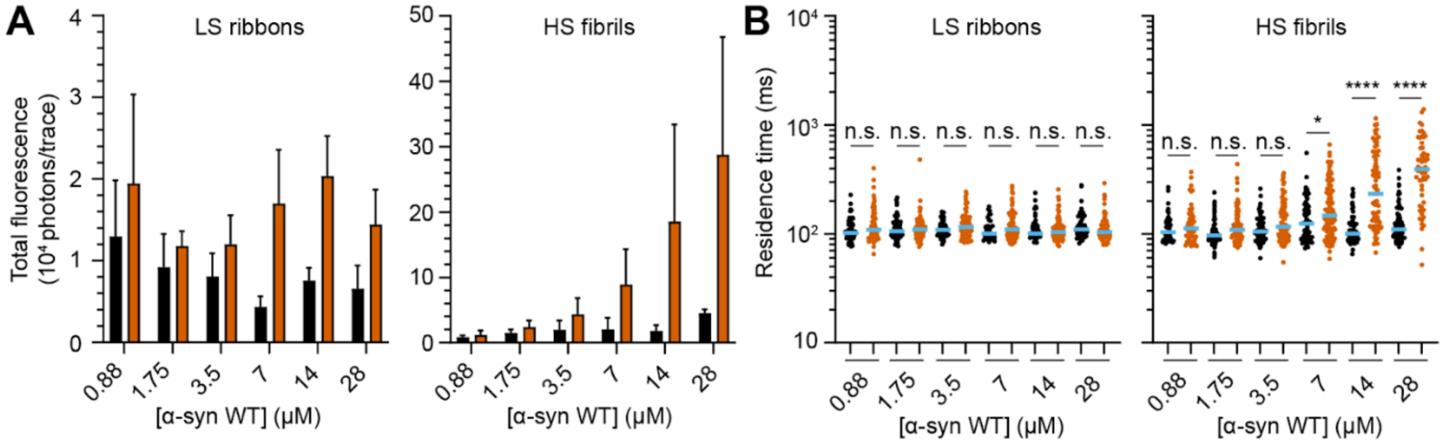


**Supporting Figure 4. Effect the growth of recombinant α-syn strains as a function of α-syn monomers.**

SmSAA was performed in a high salt buffer as a function of monomeric α-syn WT. **A.** Bar graph plotting the total fluorescence intensity of seeded reactions before (black) and after amplification. Error bar is standard deviation. **B.** Logarithmic scatter plot comparing the residence time of detected peaks in seeded reaction before (black) and after amplification (vermillion). The median value is highlighted in blue. Each symbol represents an individual event and statistics used Kruskall-Wallis test, $p \leq 0.0001$ (****), $p \leq 0.05$ (*), nonsignificant (n.s.).

**Supporting Figure 5**

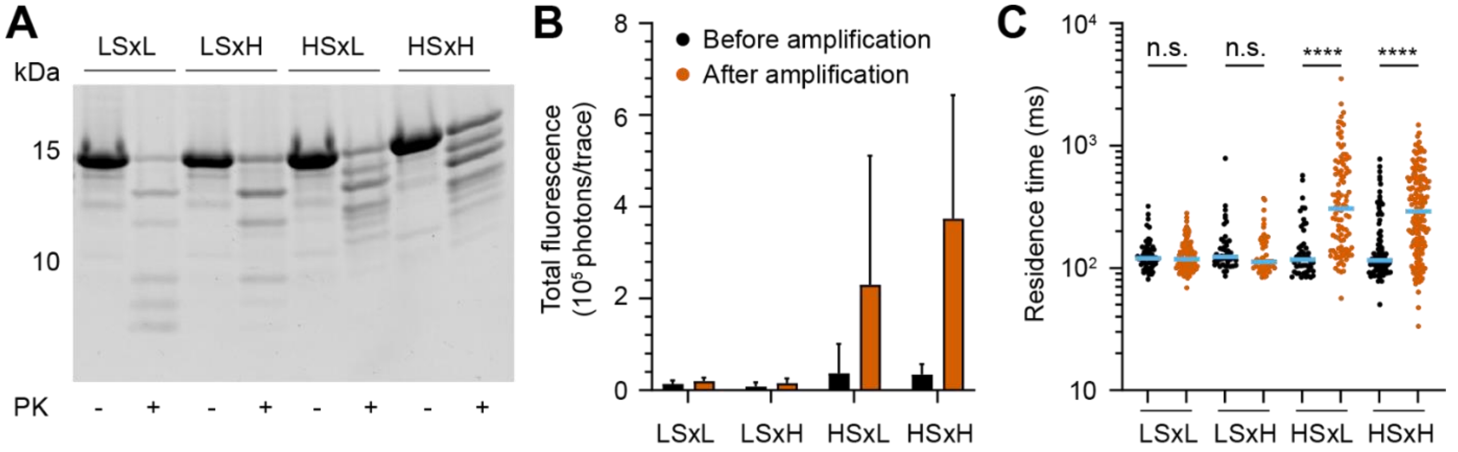


**Supporting Figure 5. Validation of cross-seeded aggregates used for cell studies.**

**A.** Digestion profile using PK for 15 min of seeded ribbons and fibrils seeded and cross-seeded in low salt buffer (LSB) or high salt buffer (HSB) for cell study. **B-C.** SmSAA phenotyping of seeded α-syn aggregates. Bar graph plotting the total fluorescence intensity of seeded reactions before (black) and after amplification with α-syn K23Q (vermillion). Error bar is standard deviation. Logarithmic scatter plot comparing the residence time of detected peaks in seeded reaction before and after amplification with α-syn K23Q. The median value is highlighted in blue. Each symbol represents an individual event and statistics used Kruskall-Wallis test, $p \leq 0.0001$ (****), nonsignificant (n.s.).

**Supporting Figure 6**

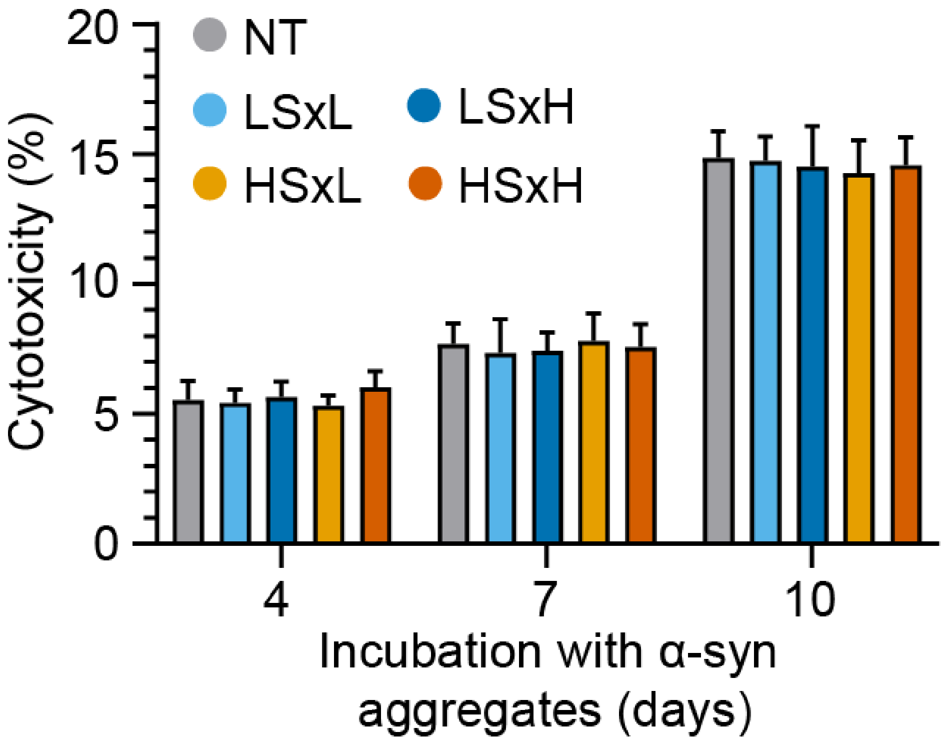


**Supporting Figure 6. Cytotoxicity of cross-seeded aggregates in SH-SY5Y cells.**

Assessment of lactate dehydrogenase levels in SH-SY5Y cells post-treatment with 5 µg/mL cross-seed α-syn aggregates: low salt ribbons seeded in low salt buffer (LSxL, sky blue), low salt ribbons seeded in high salt (LSxH, blue), high salt fibrils seeded in low salt buffer (HSxL, orange) and high salt fibrils in high salt buffer (HSxH, vermillion) over time via colorimetric cytotoxicity assay. The outcomes are represented as the percentage of cytotoxicity in maximum lactate dehydrogenase release. Error bars represent mean ± standard deviation.

**Supporting Figure 7**

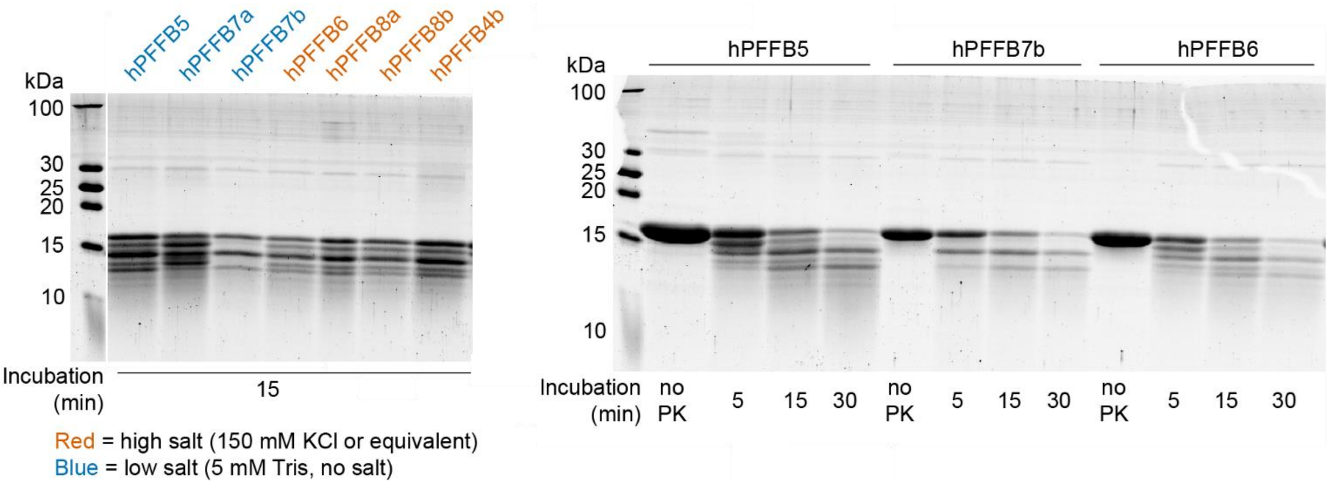


**Supporting Figure 7. Variation of polymorphs in the preparation of some Low Salt aggregates.**

When we generated the different batches of ribbons, we noticed that the PK digestion patterns of some batches deviated from the expected result (typical shown in hPFFB7b). Variations were even observed from the same master mix, separated into two tubes (Batch 7a and 7b – hPFFB7a/b). The reactions occurred in the exact same conditions of shaking and temperature. We did not observe the same variability in the fibrils generated in high salt conditions.

**Supporting Figure 8**

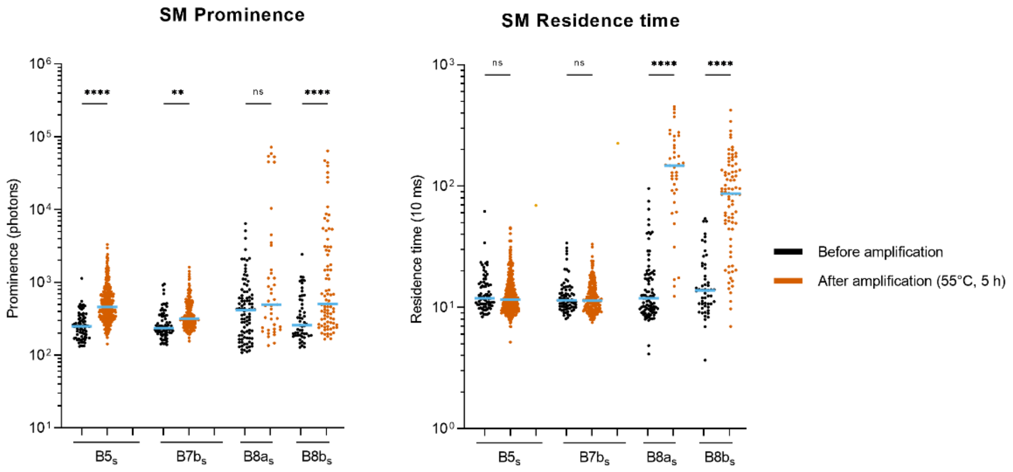


**Supporting Figure 8. Amplification of different batches of ribbons and fibrils**

Despite apparent differences in PK digestion patterns, the different ribbons obtained all displayed the same propensity for amplification, with a small increase in residence time as compared to fibrils.

### Mathematical modelling:

To obtain the equations that describe aggregate growth in RT-QuIC experiments, the approach developed by Knowles et al. was followed[3].

We describe polymers with two numbers : their number of monomers $(i)$ and their number of growth points $(j)$. For example, if the polymer is linear, $j = 2$. If it has three branches, $j = 5$.

$f(t, i, j)$ is the concentration of polymers that have these properties. We will then be interested in the polymer mass concentration $M(t) = \sum_{i,j} if(t, i, j)$ and the concentration of growth points $N(t) = \sum_{i,j} jf(t, i, j)$. We also introduce the concentration of free monomers $m(t)$.

We consider three elementary reactions in this model : **elongation** - adding a monomer on a growth point, **secondary nucleation** - adding a new growth point in the bulk of a polymer, and **fragmentation** - breaking a polymer in two and creating two new growth points. $M, N, m$ and the elementary reactions are depicted in figure 5 in the main text.

We consider situations without primary nucleation in this work, as we discuss the effect of different processes in seed amplification assays, where conditions are optimized to suppress formation of aggregates in non-seeded samples.

The smallest polymer size is considered to be 2. At $t = 0$, linear polymer seeds $(j = 2)$ are added in a solution of free monomers. The total concentration of monomers is $m_{tot}$. The initial polymer mass concentration is $M_0$, and the initial growth points concentration is $N_0$.

We first consider a situation where there is only elongation. The number of growth points per polymer is always 2. We obtain the following equations:

$$\frac{\partial f(t, i)}{\partial t} = 2k_e m(t)[f(t, i-1) - f(t, i)]$$

where $k_e$ is the elongation rate and $m(t) = m_{tot} - M(t)$. Hence:

$$\frac{dN}{dt} = 0$$

$$\frac{dM}{dt} = 2k_e m(t) \sum_{i=2}^{+\infty} i[f(t, i-1) - f(t, i)]$$

$$= k_e m(t) \sum_{i=2}^{+\infty} 2f(t, i)$$

$$= k_e m(t) N(t) = k_e m(t) N_0$$

As $N$ is a constant, we only have one differential equation that we can solve for a set of initial conditions :

$$M(t) = m_{tot} - (m_{tot} - M_0)e^{-k_e N_0 t}$$

$$\Rightarrow m(t) = (m_{tot} - M_0)e^{-k_e N_0 t}$$

If we add fragmentation, we obtain the equations developed in the publication by Knowles et. al[3].

We still have $j = 2$ for all polymers.

$$\frac{\partial f(t,i)}{\partial t} = 2k_e m(t)[f(t,i-1) - f(t,i)] - k_f(i-1)f(t,i) + 2k_f \sum_{p=i+1}^{+\infty} f(t,p)$$

where $k_f$ is the fragmentation rate.

This leads to the following set of coupled equations:

$$\frac{dN}{dt} = k_f[2M(t) - 3N(t)]$$

$$\frac{dM}{dt} = \left[m(t)k_e - k_f\right]N(t)$$

In the case of secondary nucleation, the number of growth points $j$ is not imposed.

We get:

$$\frac{\partial f(t,i,j)}{\partial t} = jk_e m(t)[f(t,i-1,j) - f(t,i,j)] + k_s(i-j)[f(t,i-1,j-1) - f(t,i,j)]$$

where $k_s$ is the secondary nucleation rate.

We consider that secondary nucleation cannot happen on the ends of the polymer, hence the $(i-j)$ factor in the above equation. We obtain:

$$\frac{dN}{dt} = \sum_{j=2}^{\infty} j \sum_{i=2}^{\infty} jk_e\, m(t)[f(t,i-1,j) - f(t,i,j)] + k_s m(t)(i-j)[f(t,i-1,j-1) - f(t,i,j)]$$

$$\frac{dN}{dt} = k_s m(t) \sum_{i,j} (i-j) f(t,i,j)$$

$$\frac{dN}{dt} = k_s m(t)[M(t) - N(t)]$$

We can do the same for $M$ and we get the following set of coupled equations:

$$\frac{dN}{dt} = k_s m(t)[M(t) - N(t)]$$

$$\frac{dM}{dt} = k_e m(t)N(t) + k_s m(t)[M(t) - N(t)]$$

Let us assume $k_s \ll k_e$ and $N \ll M$. We obtain the following approximate equations:

$$\frac{dN}{dt} = k_s m(t)M(t)$$

$$\frac{dM}{dt} = k_e m(t)N(t)$$

The following functions are solutions of this system:

$$M(t) = \frac{m_{tot}}{1 + \frac{mtot - M_0}{M_0} e^{-\kappa t}}$$

$$N(t) = \lambda M(t)$$

Where $\lambda = \sqrt{k_s/k_e}$ and $\kappa = m_{tot}\sqrt{k_e k_s}$

We can combine fragmentation and secondary nucleation. We would need other terms to account for the specifics of fragmentation on branched polymers. For example, if fragmentation occurs at a secondary nucleation site, the effect on $N$ is different, as only one new growth point is created. To simplify, we used the previous approximation $k_f \ll k_e$ and the equations become:

$$\frac{dN}{dt} = k_s m(t)M(t) + 2k_f M(t)$$

$$\frac{dM}{dt} = k_e m(t)N(t)$$

We can also find these macroscopic equations by considering the effects of the three elementary reactions on $M$ and $N$.

**Fitting of experimental curves:**

In RT-QuIC experiments, the dye ThT is a marker of amyloid formation, so the fluorescent signal obtained is proportional to the polymer mass concentration (M(t) in our model). We observe that fibrils seem to amplify much faster than ribbons, and we want to know if this difference can be explained by secondary nucleation. In order to do this, we numerically generate curves for M(t) and fit the experimental data by varying the parameters. According to equation 3 we have 6 relevant parameters : $M_0$, $N_0$, $m_{tot}$, $k_e$, $k_s$, $k_f$ . We cannot reliably fit 6 parameters based on a curve with two inflexion points. Moreover, the equations show that fragmentation and secondary nucleation have very similar effects on M(t). Indeed, both phenomena increase the value of N, which in turn increases the growth rate of M. Consequently, it will be difficult to fit $k_s$ and $k_f$ independently.

The total amount of monomers is the same for all experiments, so we can use it as a scale for all concentrations and set $m_{tot}$ = 1. This means that $m_{tot}$ becomes our concentration unit. The elongation rate should be similar for all experiments, so we arbitrarily set its value to a constant $k_e = 1000\, m_{tot}^{-1}.s^{-1}$. If one considers that the typical size of a ribbon/fibril seed is approximatively a thousand monomers, we will fit $M_0$ and estimate that $N_0$=$M_0$/500.

As $k_s$ and $k_f$ have similar effects on the evolution of M(t), we set $k_s$=0 and fitted the curves to obtain an artificial $k^*_f$ that takes into account both fragmentation and secondary nucleation.

As one can estimate that the real fragmentation rate is likely to be similar in all experiments, as it is caused by quaking, we can assume that major differences in $k^*_f$ will reflect changes in secondary nucleation.

To sum up, the values of $m_{tot}$,$\frac{M_0}{N_0}$, $k_e$ and $k_s$ were fixed, and the fit is simply performed by adjusting $M_0$ and $k^*_f$.

There are 6 experimental curves, 3 obtained from ribbon strains and 3 obtained from fibril strains. The fragmentation/secondary nucleation rate $k^*_f$ is 5.65.10$^{-4}$.s$^{-1}$ for fibrils and 5.31.10$^{-4}$ s$^{-1}$ for ribbons.

If we assume that the fragmentation rates are of the same magnitude for both strains, we can conclude that in the case of fibrils, secondary nucleation is much more important than fragmentation.

If we consider that the fragmentation rate $k_f$ for the fibrils is the same as the ribbons, ie $k^*_f$ obtained previously, then we obtain:

$$k_f^* \approx k_f + \frac{k_s}{2} m_{tot}$$

Leading to $k_s \approx 200 k_f / m_{tot} \sim 10^{-3}\ s^{-1}\ m_{tot}^{-1}$

As we set the elongation rate $k_e = 1000\ s^{-1}\ m_{tot}^{-1}$ we obtain $\lambda = \sqrt{k_s/k_e} \sim 10^3$

And as $N(t) \sim \lambda M(t)$, we deduce that one branching point (or secondary nucleation event) is formed every 2,000 monomers along the growing fibril.

**References**


1. Lau, D.; Magnan, C.; Hill, K.; Cooper, A.; Gambin, Y.; Sierecki, E., Single Molecule Fingerprinting Reveals Different Amplification Properties of α-Synuclein Oligomers and Preformed Fibrils in Seeding Assay. *ACS chemical neuroscience* **2022,** *13* (7), 883-896.

2. Bhumkar, A.; Magnan, C.; Lau, D.; Jun, E. S. W.; Dzamko, N.; Gambin, Y.; Sierecki, E., Single-Molecule Counting Coupled to Rapid Amplification Enables Detection of α-Synuclein Aggregates in Cerebrospinal Fluid of Parkinson's Disease Patients. *Angewandte Chemie (International ed. in English)* **2021,** *60* (21), 11874-11883.

3. Knowles, T. P.; Waudby, C. A.; Devlin, G. L.; Cohen, S. I.; Aguzzi, A.; Vendruscolo, M.; Terentjev, E. M.; Welland, M. E.; Dobson, C. M., An analytical solution to the kinetics of breakable filament assembly. *Science (New York, N.Y.)* **2009,** *326* (5959), 1533-7.
